# A Single-Cell Atlas and Lineage Analysis of the Adult Drosophila Ovary

**DOI:** 10.1101/798223

**Authors:** Katja Rust, Lauren Byrnes, Kevin Shengyang Yu, Jason S. Park, Julie B. Sneddon, Aaron D. Tward, Todd G. Nystul

## Abstract

The Drosophila ovary is a widely used model for germ cell and somatic tissue biology. We have used single-cell RNA-sequencing to build a comprehensive cell atlas of the adult Drosophila ovary containing unique transcriptional profiles for every major cell type in the ovary, including the germline and follicle stem cells. Using this atlas we identify novel tools for identification and manipulation of known and novel cell types and perform lineage tracing to test cellular relationships of previously unknown cell types. By this we discovered a new form of cellular plasticity in which inner germarial sheath cells convert to follicle stem cells in response to starvation.

**Graphical Abstract:** 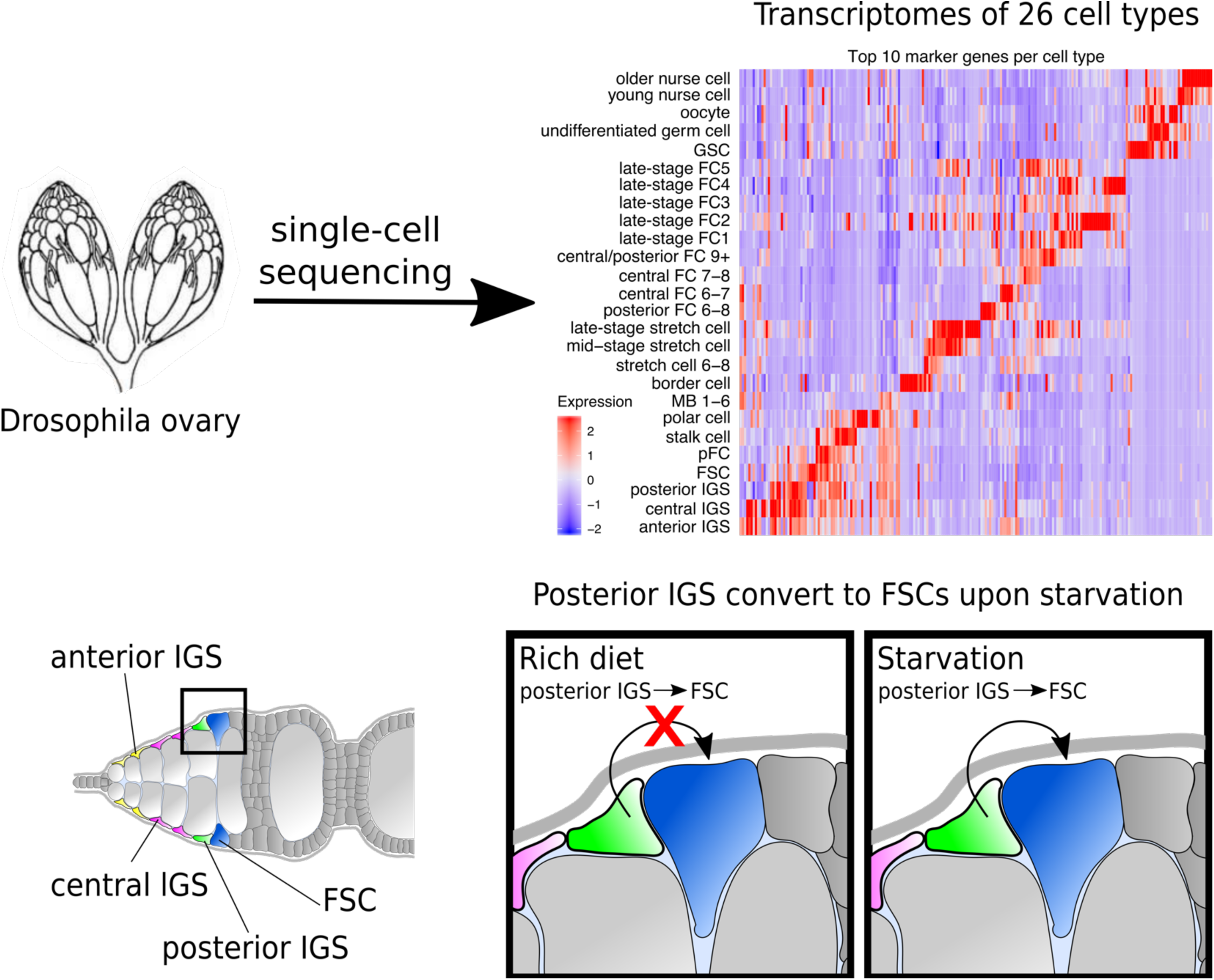

## Introduction

Decades of research on the ovary have built a foundation of knowledge that make this organ a highly tractable experimental model for studies of well-conserved biological processes, including meiosis, oogenesis, epithelial morphogenesis, and stem cell biology.^1–5^ However, our understanding of the ovary is limited by the tools available, and strategies to identify new genetic tools have been incomplete. In this study, we combined single-cell RNA-sequencing (scRNA-Seq) with methods for *in vivo* validation to develop a cell atlas of the *Drosophila* ovary that describes the identity and location of 26 bioinformatically distinct populations and the lineage relationships between key somatic cell types. We identified transcriptional profiles for various known cell types that could not previously be isolated from wildtype tissue, such as germline stem cells (GSC), follicle stem cells (FSCs), and prefollicle cells (pFCS) and describe a transcriptional timeline in main body follicle cells. In addition, we provide new evidence for transcriptionally distinct populations of IGS cells, nurse cells, and main body follicle cells. Using newly identified tools, we found that the posterior IGS cells are functionally distinct and can convert to FSCs upon starvation. This reveals an unexpected form of cellular plasticity that may be important for maintaining tissue homeostasis in response to physiological stress. Collectively, our study provides a new resource for the use of the ovary as an experimental model and demonstrates the utility of this resource for identifying and characterizing new populations of cells in the ovary.

## Results

### Identification of distinct cell types in the ovary

Each ovary is composed of approximately 16 strands of developing follicles, called ovarioles, and oogenesis begins at the anterior tip of each ovariole in a structure called the germarium (Fig. 1A-B). Two to three GSCs reside in a niche at the anterior edge of the germarium and divide during adulthood to self-renew and produce cystoblasts that move toward the posterior as they progress through oogenesis.^6^ In addition, cap and terminal filament cells provide a niche for the GSCs; IGS cells promote early germ cell differentiation; and the FSC lineage facilitates the formation of the germ cell cyst into a follicle that buds off from the germarium. Germ cells undergo four rounds of incomplete mitosis to form into a cyst of 16 interconnected cells. One cell is selected to become the oocyte and enters meiosis, while the other 15 become polyploid “nurse” cells that provide nutrients, organelles, and macromolecules to the oocyte. At the midpoint in the germarium, each cyst becomes encapsulated by a layer of epithelial follicle cells produced by the FSCs.^7^ FSCs divide with asymmetric outcomes to self-renew and produce pFCs that differentiate gradually, over the course of several divisions ^8^ into polar cells, stalk cells, or main body follicle cells (Fig. 1A). Under ideal conditions, newly budded follicles grow and develop into a mature egg over 4-5 days.^9, 10^ This stereotypical process has been divided into 14 distinct stages, with early stages (Stages 1-6) characterized by rapid follicle growth and follicle cell division; mid-stages (Stages 7-10) characterized by the onset of yolk protein production, elongation of the follicle, growth of the oocyte, and specialization of follicle cells into subtypes such as stretch cells; and late stages (Stages 11-14) characterized by the death of nurse cells, deposition of the egg shell proteins, and growth of the oocyte to fill the entire volume inside the egg shell (Fig. 1B).

**Figure 1:**
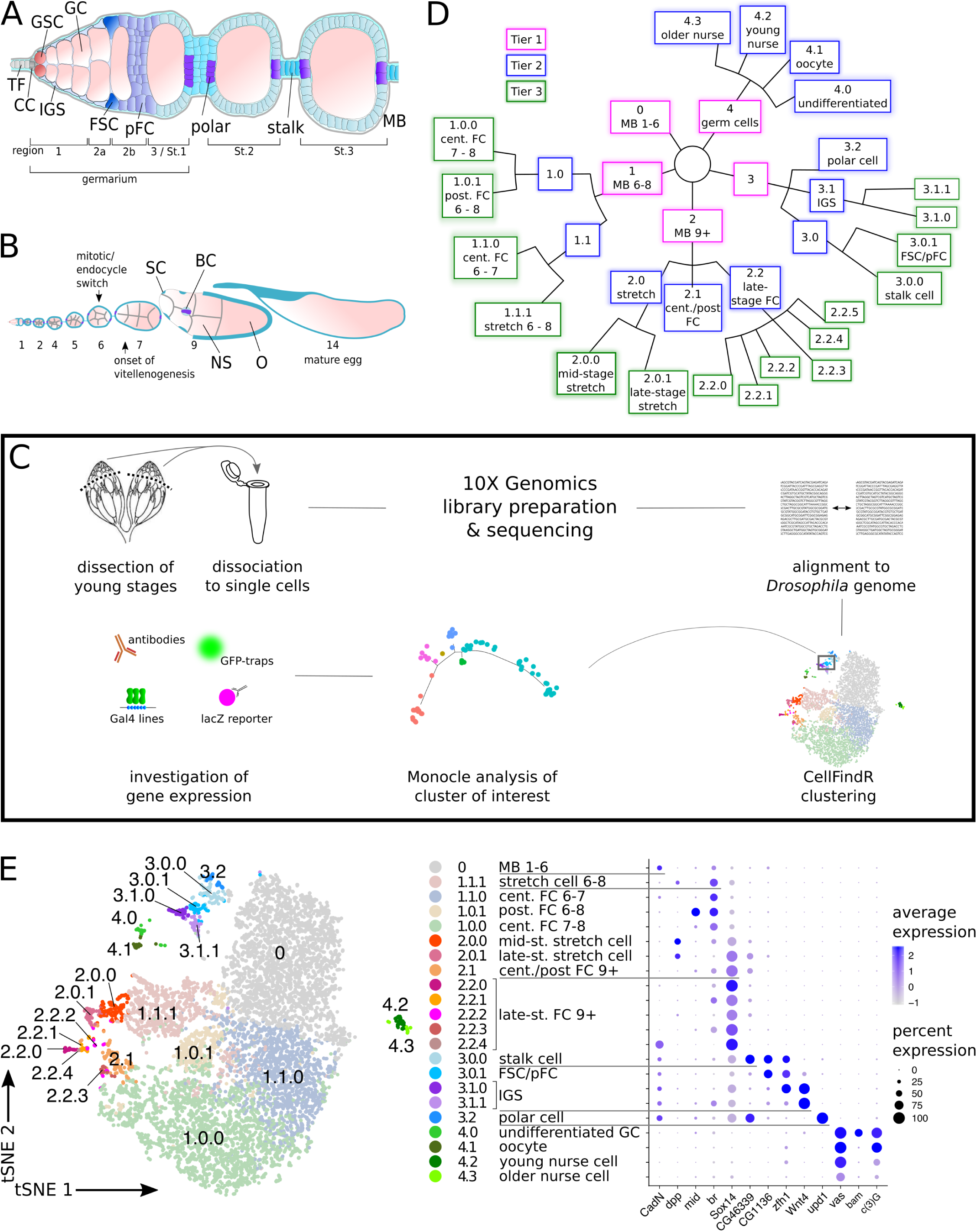
CellFindR identifies distinct populations of cells in the ovary. A) Diagram of the anterior tip of the ovariole, including the germarium and two budded follicles. B) Diagram of the entire ovariole. C) Schematic overview of sample preparation for single-cell sequencing. D) Hierarchy of CellFindR clusters. Tier 1 (magenta), Tier 2 (blue) and Tier 3 (green) clusters were produced by the first, second and third round of CellFindR clustering, respectively. E) CellFindR tSNE plot of 9623 cells isolated from Drosophila ovaries and gene expression profiles of selected markers. St: Stage; MB: main body follicle cells; FC: follicle cells; FSC: follicle stem cells; IGS: inner germarial sheath cells; GC: germ cells.

To catalog the cell types in the *Drosophila* ovary, we dissected ovarioles from over 60 wildtype flies, dissociated the tissue to a single-cell suspension, and performed scRNA-Seq (Fig. 1C). This procedure produced transcriptional profiles of approximately 10,000 cells, achieving about 1.5x coverage of the ovariole (see Materials and Methods). We clustered the cells using an adaptation of the Seurat algorithm^11, 12^ called CellFindR that was first described in a study of mammalian cochlear development ^13^. Seurat organizes the cells in a dataset into clusters of cells that have similar transcriptional profiles, with the number of clusters determined by a user-specified resolution factor. However, in many cases, it is not possible to accurately cluster the entire dataset with a single iteration of the Seurat algorithm. Therefore, a common strategy is either to over-cluster the dataset with a very high resolution factor and then manually merge back clusters that appear to be related, or to under-cluster the dataset and then re-run Seurat on a subset of clusters that appear to contain biologically relevant heterogeneity. CellFindR not only automates these steps but also performs them in a structured and unbiased manner. Specifically, CellFindR, first applies the Seurat algorithm to the entire dataset, producing a set of “Tier 1” clusters, and then applies it to each Tier 1 cluster separately to test whether they can be further sub-clustered. Newly produced clusters are designated as “Tier 2” clusters, and this process is then repeated, producing new tiers with each iteration, until no additional clusters can be identified. We found that this method was more accurate at producing clusters that aligned with markers of known cell types than using Seurat alone (Fig. S1A-M). In addition, CellFindR produces clusters in a way that allows them to be arranged in a hierarchical tree, with major branches separating distantly related cell types and more minor branches separating more closely related cell types (Fig. 1D). The hierarchical tree outperformed clustering methods based on average gene expression (Fig. S1N).

When applied with standard settings, CellFindR parsed the cells into five Tier 1 clusters, including one germ cell cluster and four somatic cell clusters that correspond broadly to different stages of oogenesis. Specifically, Cluster 3 contains somatic cells in the germarium, polar cells, and stalk cells; Cluster 0 contains the somatic cells in early-stage follicles; Cluster 1 contains the somatic cells in mid-stage follicles; Cluster 2 contains the somatic cells in late-stage follicles; and Cluster 4 contains germ cells. In Tier 2, CellFindR parsed the germ cell cluster (Cluster 4) into four clusters, two of the somatic cell clusters (Clusters 2 and 3) into three clusters each, and left the remaining somatic cell clusters (Cluster 0 and 1) intact. With a higher resolution setting, CellFindR parsed Cluster 1 into four Tier 2 clusters. Lastly, CellFindR parsed several Tier 2 clusters into Tier 3 clusters, producing a total of 22 bioinformatically distinct clusters (Fig. 1C-D, Tables S1-2). Notably, the predicted cellular identity and transcriptional profiles of these clusters align closely with a second scRNA-Seq dataset that we generated independently (Fig. S2, Table S3).

### Germ cell clustering identifies distinct populations of undifferentiated germ cells, oocytes, and nurse cells

The germ cells clustered apart from somatic cells into a single Tier 1 cluster (Cluster 4) and are distinguished by the expression of germ cell markers such as *vasa* (*vas*)^14, 15^ and the lack of expression of somatic cell markers such as *traffic jam* (*tj*) (Fig. 2A-B)^16^. The four Tier 2 germ cell clusters can be distinguished from each other by specific combinations of marker genes (Fig. 2C-G). Cluster 4.0 is enriched for cells that express genes such as *bam* and *bgcn*, indicating that this cluster corresponds to the early, undifferentiated stages of germ cell development (Fig. 2D, H)^17–19^. To test this assignment experimentally, we determined the expression pattern of *groucho* (*gro*), which is a novel predicted marker of Cluster 4.0 (Fig. 2H, S3A). Indeed, we found that *gro* is strongly expressed in Region 1 and 2a germ cell cysts in wildtype ovaries, which is where *bam* and *bgcn* are expressed, and then tapers off in more mature cysts (Fig. 2O). Cluster 4.1 is enriched for cells that express genes involved in the formation and function of the synaptonemal complex, such as *corona* (*cona*), *c(3)G*, and *c(2)M*^20^ (Fig. 2H, Fig. S3B-D), indicating that this cluster contains oocytes. Interestingly, the transcripts of these genes are also expressed in some cells in Cluster 4.0, which corresponds to the early stages of germ cell development (Fig. 2H, Fig. S3B-D). This is consistent with the observation that synaptonemal complexes begin to form in Region 2a cysts, and can be observed in both the future oocyte and other germ cells in the cyst at this stage^21^. Lastly, the cells in Clusters 4.2 and 4.3 have little or no meiotic gene expression but are distinguished from each other by the expression of genes such as *jtb* in Cluster 4.2 and *26-29-p* in Cluster 4.3 (Fig. 2F-H, S3B-D). Both clusters are enriched for cells with increased expression of genes such as *CycE* and *ago* (Fig. 2H, Fig. S3E-F), which promote endocycling^22^, indicating that they correspond to the nurse cell populations.

**Figure 2:**
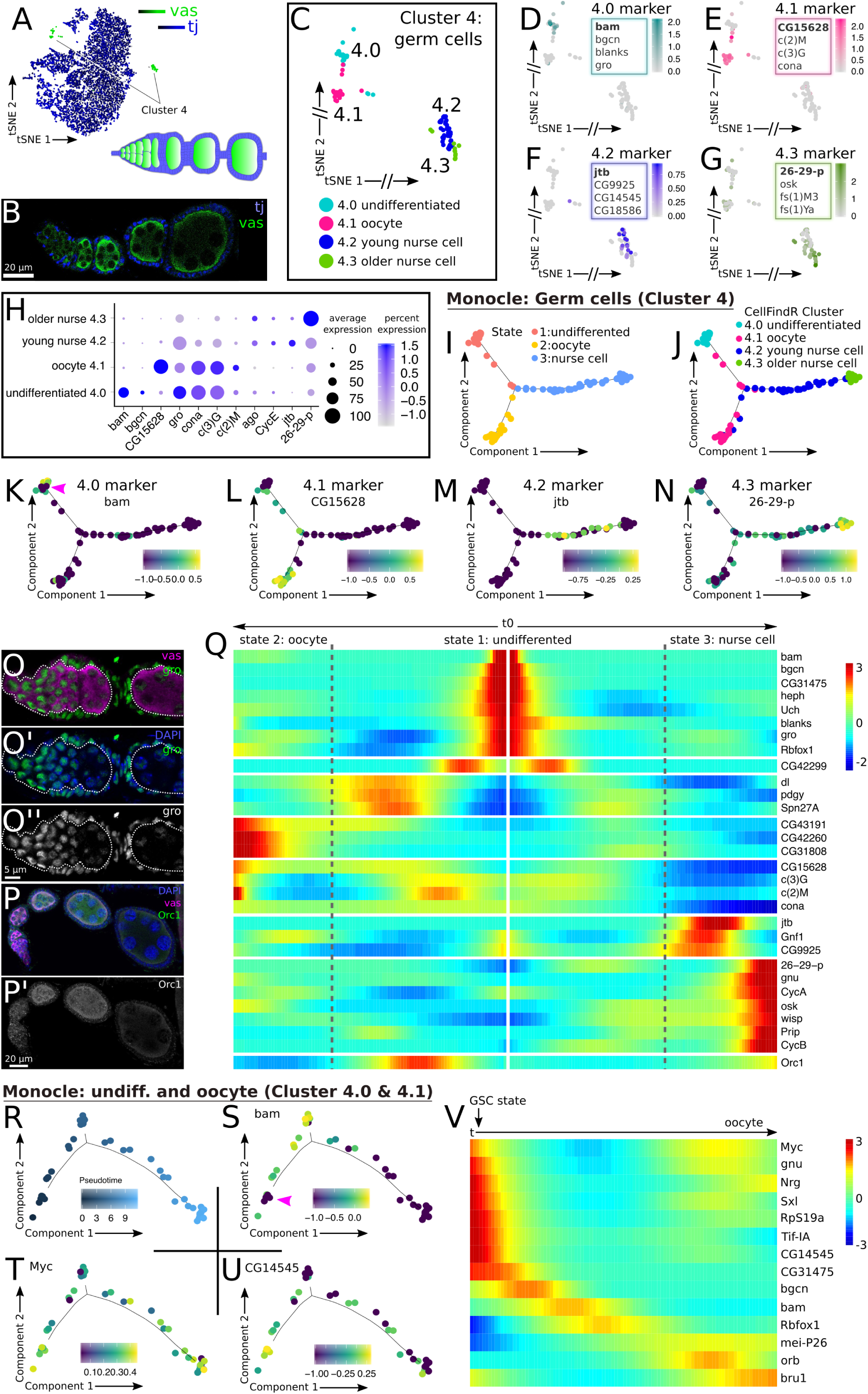
Germ cells. A) SCope expression plot of *vas* (green) and *tj* (blue) on tSNE projection of CellFindR plot and a diagram of an ovariole showing cell types in the corresponding colors. B) Early stages of Drosophila ovariole stained with tj (blue) and vas (green). C) tSNE plot of CellFindR clusters containing germ cells. D-G) Germ cell markers. Expression of the marker in bold text is shown. H) Expression profile of selected markers in germ cell clusters. I-J) Monocle analysis of germ cells identifies three different states in pseudotime (I) that correspond to the four stages of differentiation identified by CellFindR (J). K-N) Expression of selected genes in pseudotime. Pink arrowhead points out *bam*^-^ cells at the beginning of pseudotime. O) Wildtype germarium stained for vas (magenta), gro (green), and DAPI (blue). gro is highly expressed in germ cells until region 2a and is expressed in all somatic cells. P) Wildtype germarium stained for vas (magenta), Orc1 (green), and DAPI (blue). Orc1 is expressed at higher levels in the germ cells of Stage 2-5 follicles. Q) Heatmap of gene expression in pseudotime of all germ cells, with the tip of State 1 in the center and the bifurcations leading to States 2 and 3 arrayed outward to the left and right, respectively. R-U) Monocle trajectories of Clusters 4.0 and 4.1 pseudocolored to indicate pseudotime (R), or expression of the indicated gene (S-U). Cells at the beginning of pseudotime express low levels of *bam* (pink arrowhead in S). V) Heatmap of gene expression in pseudotime of germ cells of clusters 4.0 and 4.1 with cells arrayed from left to right according to pseudotime. The earliest time point marks the germ line stem cell (GSC) state.

To estimate the lineage relationships among the germ cells in our dataset, we performed Monocle analysis on the entire population in Cluster 4. Monocle is an algorithm that arranges cells along a bioinformatic trajectory that minimizes the differences in gene expression between neighboring cells^23, 24^. When applied to a set of cells in the same lineage, the cells are organized in “pseudotime” according to the stage of differentiation. Monocle arranged the cells in Cluster 4 into a trajectory with three branches (Fig. 2I). Germ cells progress along a common path during the early mitotic stages and then differentiate into either an oocyte or a nurse cell, so it is likely that one branch represents the early stages of differentiation and the other two represent the paths toward the oocyte and nurse cell fates. Consistent with this prediction and CellFindR clusters, the tip of one branch is enriched for cells in Cluster 4.0 expressing markers of the early stages of differentiation, such as *bam* and *bgcn*, one branch is enriched for cells expressing oocyte markers in Cluster 4.1 and meiotic genes, and the third branch is enriched for nurse cell markers in Cluster 4.2 and 4.3 (Fig. 2I-N). On the nurse cell branch, the earlier cells express Cluster 4.2 markers, such as *jtb*, whereas the later cells express Cluster 4.3 markers such as *26-29-p* (Fig. 2J,M-N), suggesting that the nurse cells in Cluster 4.2 are younger than those in Cluster 4.3. We queried for genes expressed at distinct time points during germ cell development (Table S4). The oldest nurse cells in pseudotime express *Orc1*, and we found that staining for *Orc1* by immunofluorescence first becomes detectable in the germ cells of newly budded (Stage 2-3) follicles (Fig. 2P-Q), suggesting that this is the oldest stage of germ cell development present in our dataset. It is likely that we did not capture germ cells from follicles older than this stage because they were too big (>10 μm by Stage 4-5^25^) to be enveloped by the oil droplets used to separate the cells for single-cell sequencing. A peak of *Orc1* expression is also detectable at an earlier stage, in the region that contains the undifferentiated germ cells, but we did not see expression at these stages at the protein level *in vivo* with *Orc1-GFP* (Fig. 2P-Q).

We noted a small population of cells that were negative for *bam* and *bgcn* in Cluster 4.0 and at the beginning of germ cell pseudotime, although Monocle failed to separate these cells into a distinct state when applied to all germ cells (Fig. 2I,K). To test whether these cells might correspond to germline stem cells (GSCs), we repeated the Monocle analysis specifically on the undifferentiated germ cell and the oocyte clusters (Clusters 4.0 and 4.1) in anticipation of a better resolution. Indeed, we found that the cells at the earliest stage of this pseudotime trajectory exhibited the expected GSC transcriptional profile, with low levels of *bam* and *bgcn* expression and high levels of *myc* and *sxl* expression (Fig. 2R-T, V, Table S5).^26, 27^ This analysis revealed several additional genes that are predicted to be specific for GSCs but have not yet been studied in the ovary, including *Tif-1A* and *CG14545* (Fig. 2U-V).

### Regions 1 and 2a contain distinct but closely-related populations of somatic cells

Clusters 3.1.0 and 3.1.1 are composed of cells that express markers of the somatic cells in Regions 1 and 2a of the germarium, with the majority of cells in both clusters expressing IGS cell markers, such as *failed axons* (*fax*) and *patched* (*ptc*) (Fig. 3A-C, and S4A).^28–30^ Analysis of these two clusters identified several additional markers that are predicted to be enriched in all IGS cells relative to other cells in the ovary, including *CG44325*, *5-HT2A*, and *Ten-a* (Fig. 3D), and we confirmed this prediction by assaying for GFP expression in *CG44325-GFP* (Fig. 3M). A small subset of cells in Cluster 3.1.0 express *engrailed* (*en*), suggesting that they are a group of terminal filament and cap cells (Fig. 3E, S4B). We identified two additional genes, *Tsp42Ej* and *Nrx-1*, with similar expression patterns, which provides additional evidence that these cells are indeed a separate cell population from the other cells in Cluster 3.1.0 (Fig. S4C-D).

**Figure 3:**
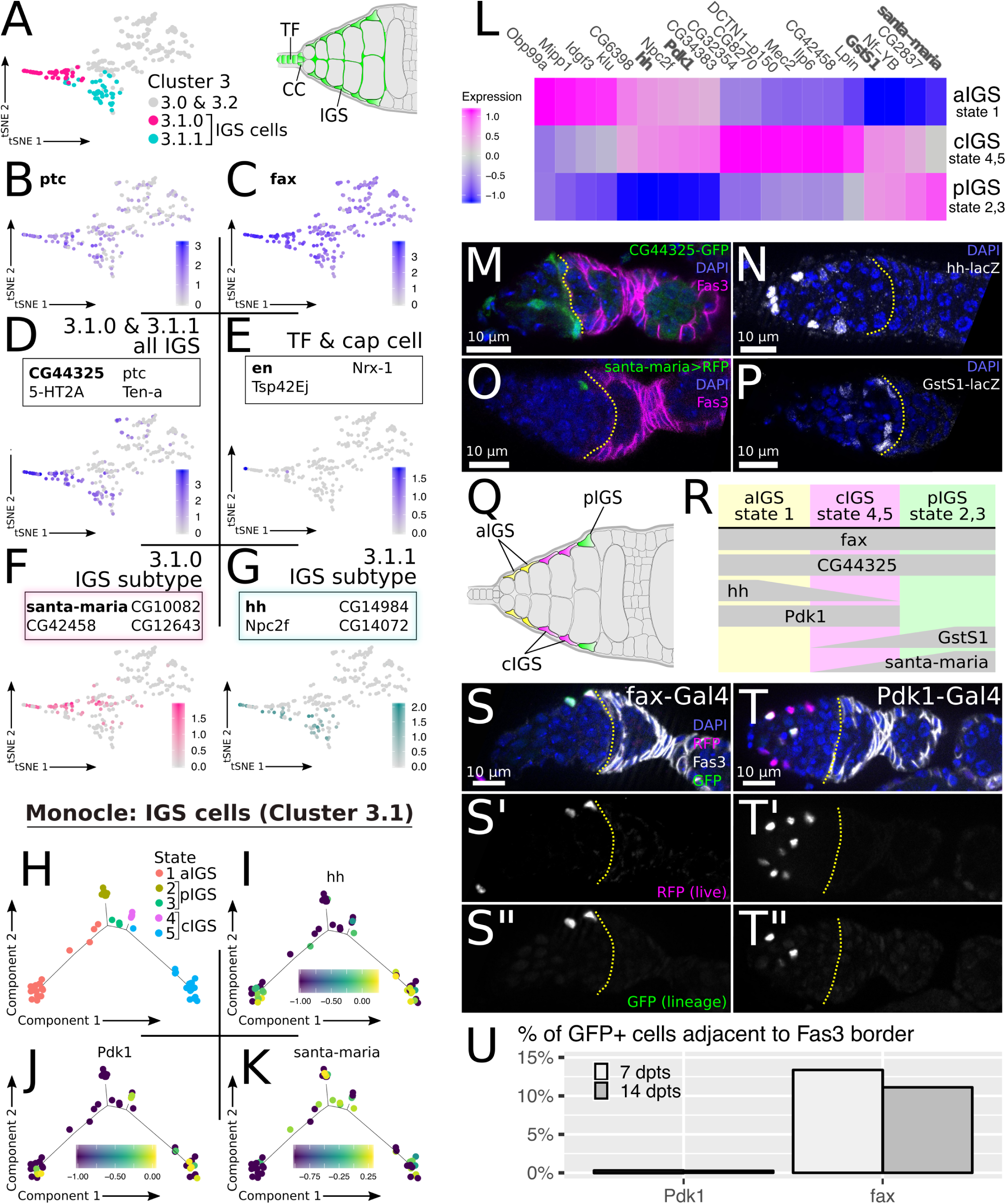
Inner germarial sheath cells. A) tSNE plot of Cluster 3. Clusters 3.1.0 and 3.1.1 contain mostly IGS cells and are highlighted. Terminal filament (TF), cap cell (CC) and IGS cells are highlighted in green on the schematic of the germarium. B-C) Expression of *ptc* and *fax* in Cluster 3. Both genes are highly expressed in Cluster 3.1.0 and 3.1.1. D-G) tSNE plots showing the expression pattern of markers in bold text. D) CG44325 marks cluster 3.1.0 and 3.1.1. E) en is expressed in 3 cells of Cluster 3.1.0. F) santa-maria is expressed in Cluster 3.1.0. G) hh is specific for Cluster 3.1.1. H) Monocle analysis of IGS cells identifies 5 states in pseudodistance. I-K) Expression of indicated genes shown in pseudodistance. L) Heatmap showing the transcriptional distinctions between States 1, 2/3, and 4/5. Bold text indicates genes that have been assayed *in vivo*. M-P) Reporters of IGS specific genes stained for DAPI (blue), and *Fas3* (magenta) and GFP (green) or LacZ (white). Yellow line marks 2a/2b border. M) CG44325-GFP is expressed in all IGS cells. N) hh-lacZ (white) is highly expressed in aIGS and expressed in lower levels in cIGS. O) santa-maria>RFP (magenta) is sparsely expressed in pIGS. P) GstS1-lacZ marks pIGS and cIGS. Q-R) Schematics showing the predicted location (Q) and expression profiles (R) of the aIGS, cIGS, and pIGS populations. S-T) Germaria with *fax-Gal4* and *Pdk1-Gal4* driving G-TRACE^ts^ stained for DAPI (blue), Fas3 (white), RFP (magenta, marks cells expressing Gal4), and GFP (green, marks descendants of Gal4 expressing cells). S) fax sparsely labels cells throughout the IGS cell population, including IGS cells at the 2b border adjacent to the first Fas3 expressing cells. T) Pdk1 marks aIGS and cIGS cells and is usually not expressed in cells at the Fas3 border. U) Quantification of GFP positive cells at the Fas3 border in *Pdk1-Gal4* and *fax-Gal4* driving G-TRACE. n=380, 865, 81, and 187 for Pdk1 7dpts, Pdk1 14dpts, fax 7dpts, and fax 14dpts, respectively.

Recent studies have identified distinct functions and morphologies for some subsets of IGS cells,^31–35^ indicating that the population is heterogeneous. In accordance with this, we identified markers specific for Cluster 3.1.0 or 3.1.1 (Fig. 3F-G). However, a systematic assessment of IGS cell heterogeneity has not been reported. Thus, to investigate the heterogeneity in the IGS cell population, we removed the cap and terminal filament cells from the two clusters containing IGS cells (3.1.0 and 3.1.1) and performed Monocle analysis on the remaining cells. Although Monocle is most commonly used to study changes in pseudotime during cellular differentiation, we reasoned that it would also be useful for identifying changes in “pseudodistance” across the IGS cell population, which exhibits an anterior-to-posterior gradient of expression of genes such as *hedgehog* (*hh*) and *ptc*^30, 36–38^ (Fig. 3N, S4A). Indeed, Monocle parsed the IGS cell population into five distinct states, and we found that they correspond to the gradient of *hh* expression, with *hh^high^* cells enriched in States 1 and 5 at the bottom of the graph, and *hh^low^* cells enriched States 2, 3 and 4 at the top (Fig. 3H-I, Table S6). We also identified three additional markers: *Pdk1*, which is upregulated in States 1 and 5, and *santa-maria*, which highly expressed in states 2 and 3 and lowly expressed in states 4 and 5 (Fig. 3J-K). Upon closer inspection, we found that the transcriptional profiles of States 2 and 3 were very similar to each other, as were the transcriptional profiles of States 4 and 5. Merging these states produced three transcriptionally distinct states: State 1, State 2/3, and State 4/5 (Fig. 3L, S4E). We found that State 2/3 is strongly enriched for cells expressing *GstS1* and *santa-maria* (Fig. 3K,L), and that both of these genes are expressed by IGS cells in Region 2a. Specifically, *GstS1-LacZ* is expressed throughout Region 2a and *santa-maria-Gal4* is expressed in only the posterior-most IGS cells, located along the Region 2a/2b border (Fig. 3O,P). In addition, cells in State 2/3 express relatively low levels of hh, and hh expression decreases toward the posterior (Fig. 3I,N). This indicates that State 2/3 corresponds to posterior IGS (pIGS) cells. Conversely, State 1 is enriched for cells expressing *hh* and *Pdk1*, which are expressed in the anterior half of the IGS cell compartment (Fig. 3P,R), whereas State 4/5 corresponds to an intermediate state, with cells that express *hh*, *Pdk1*, and *GstS1*, and only low levels of *santa-maria* (Fig. 3I-L). This suggests that State 1 corresponds to anterior IGS (aIGS) cells and State 4/5 corresponds to central IGS (cIGS) cells located in between the aIGS and pIGS cells (Fig. 3Q-R).

The lineage potential of IGS cells in Regions 1 and 2a is not fully understood and has been the subject of some recent controversy. Specifically, though IGS cells have been reported to be mitotic,^32^ it is not known whether IGS cells produced in one region of the germarium intermix with IGS cells in another part of the germarium. In addition, although most studies have considered all somatic cells in Region 2a to be IGS cells, a recent study proposed that three rings of *Fas3^−^* somatic cells in the posterior half of Region 2a are FSCs rather than IGS cells.^39^ However, we subsequently evaluated this claim and found that the anterior-most cells in large, persistent FSC clones are typically *Fas3^+^*, which strongly argues against this possibility.^40^ To further test whether somatic cells in Region 2a are capable of contributing to the FSC lineage and to test, more generally, the lineage potential of the IGS cell populations identified by our single-cell sequencing analysis, we combined *fax-Gal4* or *Pdk1-Gal4* with the lineage tracing tool, G-TRACE,^41^ in which RFP expression specifically labels the Gal4-expressing cells and GFP^+^ clones trace the lineage of the Gal4-expressing cells. To ensure that the G-TRACE tool is only activated during adulthood, we also crossed in a *tub-Gal80^ts^* construct, raised flies at 18°C, and shifted to 29°C after eclosion (referred to herein as G-TRACE^ts^). Although *fax-GFP* was reported to be expressed in all IGS cells,^28, 29^ we found that *fax-Gal4*, which is inserted into an intron of *fax*, drove expression of RFP in an average of just 4.4 IGS cells positioned sporadically throughout Regions 1 and 2a, but never in *Fas3^+^* cells (Fig. 3S). After 7 or 14 days post temperature shift (dpts), we observed small GFP^+^ clones throughout the IGS cell compartment (an average of 2.6 GFP^+^ cells per germarium, n=73) in Regions 1 and 2a, but not in the FSC lineage in nearly every case (Fig. 3S, S4F). The only exceptions to this pattern were two ovarioles with FSC clones and two ovarioles with follicle cell clones, and all four of these ovarioles were isolated from the same fly at 7 dpts. This interesting outlier is considered further in Figure 7.

With *Pdk1-Gal4* driving G-TRACE^ts^ at 7dpts, we observed strong RFP expression in all IGS cells throughout the anterior and central regions of the IGS cell compartment but almost never in posterior IGS cells (0.08%, n=1283 RFP^+^ cells, Fig. 3T). The GFP^+^ clones produced by *Pdk1-Gal4* driving G-TRACE^ts^ were largely confined to this same region of the germarium and rarely expanded to the 2a region where cells did not express RFP (Fig. 3T, S4F-G). To further describe the differences in the clonal patterns in *Pdk1-Gal4* and *fax-Gal4*, we looked specifically at expression in the most posterior IGS cells, which are adjacent to the boundary of *Fas3* expression. We found that, with *Pdk1-Gal4*, GFP^+^ cells were almost never adjacent to the *Fas3* border and only 0.3% (n=865) of the GFP^+^ cells in these germaria were adjacent to the *Fas3* border (Fig. 3T-U, S4G). In contrast, with *fax-Gal4* driving G-TRACE^ts^, we frequently observed RFP^+^ cells adjacent to the *Fas3* border and 13.4% (n=187) of GFP^+^ cells in these germaria were adjacent to the *Fas3* border (Fig. 3S,U). Taken together, these data indicate the aIGS and cIGS cells that express *Pdk1-Gal4* typically do not contribute to the pIGS cell population, and pIGS cells, including those that are adjacent to the *Fas3* border, do not typically contribute to the FSC lineage.

### The early stages of differentiation in the FSC lineage exhibit distinct expression profiles

Clusters 3.0.1, 3.0.0, and 3.2 are distinguished from other clusters by the strong expression of follicle cell markers, such as *Fas3* and *Jupiter* (Fig. 4A-C, G).^42, 43^ A large majority of cells in Cluster 3.2 express *unpaired1* (*upd1*) (Fig. 4D, H, S5A), indicating that this cluster contains the polar cells.^44^ Clusters 3.0.0 and 3.0.1 (as well as 3.1.0, which contains pIGS and cIGS cells) are enriched for cells that express the transcription factor, *zfh-1*, whereas the polar cell cluster (Cluster 3.2) is not (Fig. 4F-G, S5A). To determine the identity of Clusters 3.0.0 and 3.0.1, we screened through publicly available reporter lines of genes that distinguish the two clusters. We found that *CG46339* is expressed Cluster 3.0.0 but not in Cluster 3.0.1 (Fig. 4E, S5A), and a Gal4 enhancer trap of this gene is strongly and specifically expressed in stalk cells, beginning with the first budded follicle (Fig. 4I), indicating that Cluster 3.0.0 contains stalk cells. The remaining cluster, Cluster 3.0.1, is enriched for cells that do not express either the stalk cell marker, *CG46339*, or the polar cell marker, *upd1*, but express high levels of *Jupiter* and *Fas3*, suggesting that it contains the FSCs and pFCs. We combined the markers identified in this analysis and found that it is possible to distinguish seven distinct populations of cells in the germarium and early follicles by co-staining for *zfh-1*, *Fas3*, and *Jupiter* (Fig. 4G,J).

**Figure 4:**
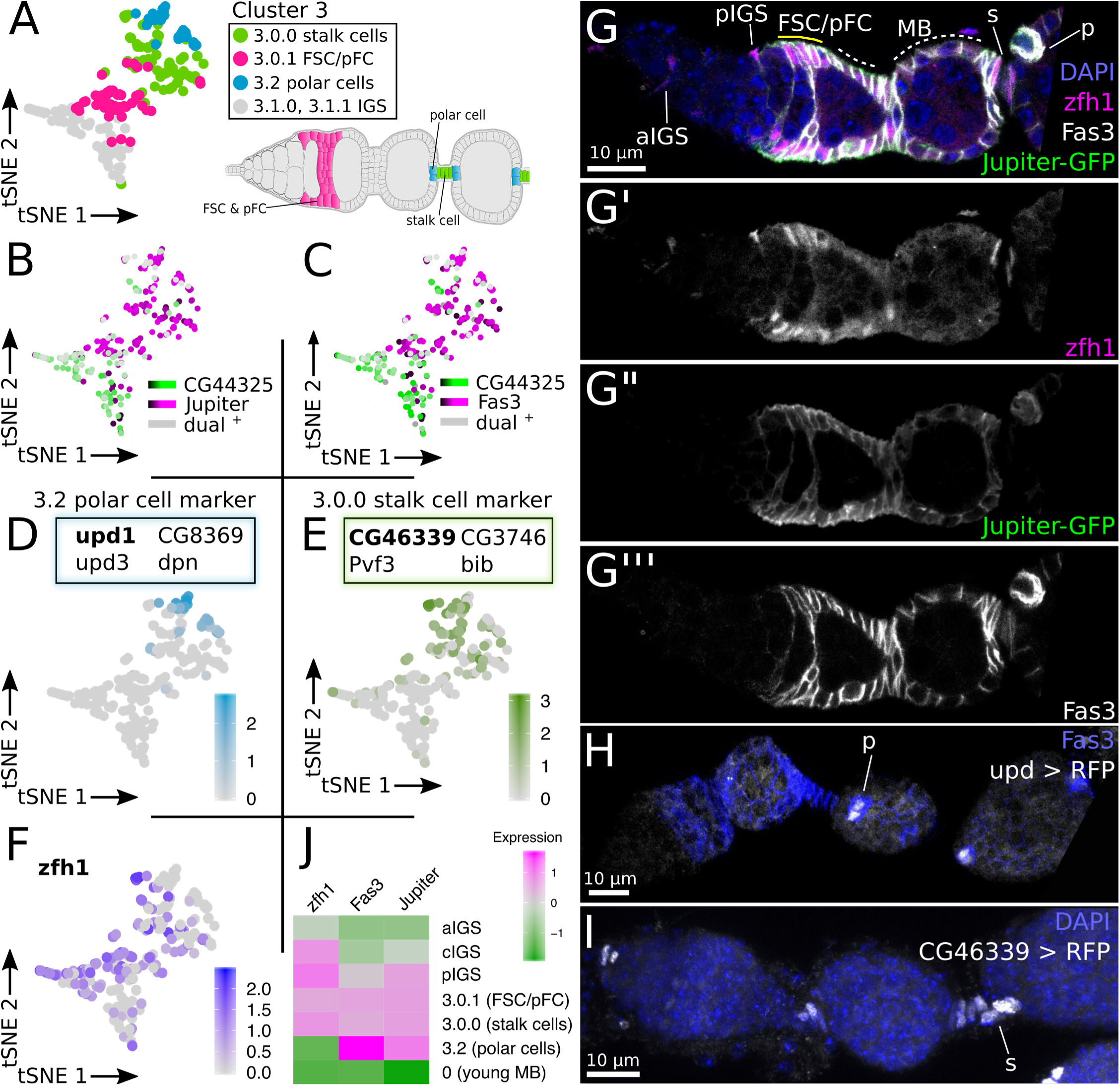
The early FSC lineage, including polar and stalk cells. A) tSNE plot of Cluster 3. Clusters 3.0.0, 3.0.1, and 3.2 primarily contain the indicated cell types, which are highlighted in the diagram of the germarium. B-C) SCope plots of Cluster 3. Main body follicle cells (Clusters 3.0.0 and 3.0.1) and polar cells are positive for *Jupiter* and *Fas3* (magenta) but do not express the IGS cell marker *CG44325* (green). D-E) Polar cell and stalk cell markers. Expression of the marker in bold text is shown. F) *zfh1* is expressed specifically in most cells in Cluster 3 (high in Clusters 3.0.0, 3.0.1, 3.1.0). G) Maximum intensity projection of upd>RFP germarium stained for Fas3 (blue) and RFP (white). RFP is specifically expressed in polar cells. H) Wildtype germarium stained for DAPI (blue), zfh1 (magenta), Fas3 (white), and Jupiter (green). This combination of staining distinguishes seven different cell populations: aIGS cells: *zfh-1^low^*, *Fas3^off^*, *Jupiter^off^*; cIGS cells: *zfh-1^high^*, *Fas3^off^*, *Jupiter^off^*; pIGS cells: *zfh-1^high^*, *Fas3^off^*, *Jupiter^high^*; FSCs and early pFCs: *zfh-1^high^*, *Fas3^high^*, *Jupiter^high^*; stalk cells: *zfh-1^high^*, *Fas3^low^*, *Jupiter^low^*, polar cells: *zfh-1^off^*, *Fas3^high^*, *Jupiter^high^*, young main body follicle cells: *zfh-1^off^*, *Fas3^medium^*, *Jupiter^medium^*. I) Maximum intensity projection of CG46339>RFP stained for DAPI (blue) and RFP (white). CG46339 is expressed in stalk cells. J) Heatmap showing the expression of *Fas3*, *Jupiter* and *zfh1*.

To obtain increased resolution into the transcriptional differences during the early stages of differentiation in the FSC lineage, we performed Monocle on the FSC/pFCs cluster (Cluster 3.0.1) and searched for genes that were significantly enriched at different points in pseudotime (Table S7). This analysis identified 10 genes that are specifically upregulated in cells at the beginning of pseudotime, prior to the upregulation of polar or stalk cell markers (Fig. 5A, S5B). These include *chickadee* (*chic*), which is known to be expressed in cells at the Region 2a/2b border,^42, 45^ and several novel markers. One of these novel markers is *GstS1* which, as we describe above, is also a marker for cIGS and pIGS cells. In the FSC lineage, we found that *GstS1* is expressed in *Fas3^+^* cells near the Region 2a/2b border but not in pFCs located further to the posterior (Fig. S5C). This confirms that these early stages of pseudotime identified by Monocle correspond to the earliest stages of the FSC lineage.

**Figure 5:**
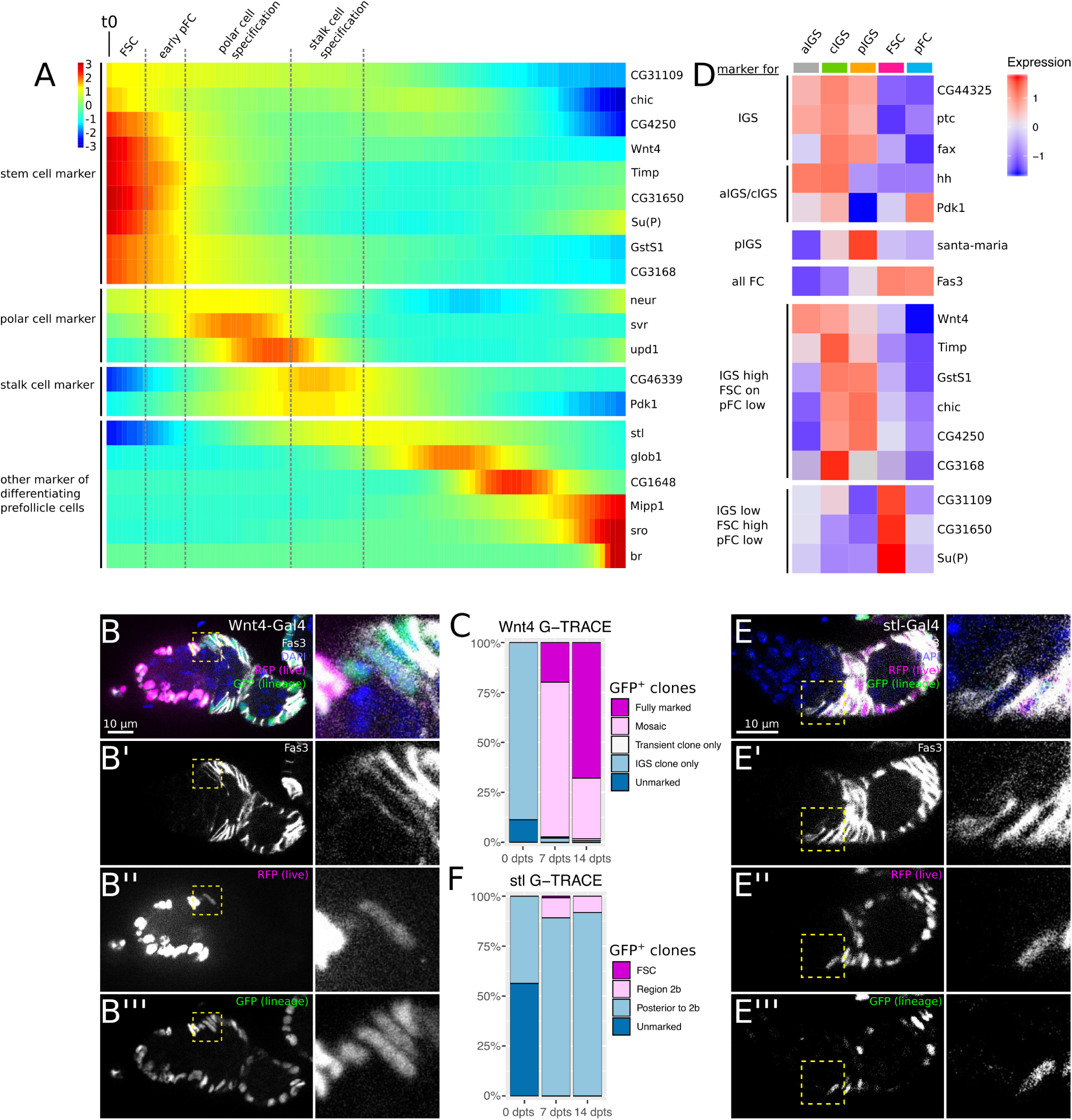
Wnt4-Gal4 specifically identifies FSCs. A) Heatmap of gene expression in pseudotime of FSCs and pFCs in Cluster 3.0.1. The earliest stages express FSC markers while older stages express markers for differentiating pFCs. See Fig. S5B for corresponding Monocle plot. B) Immunostaining of a germarium with *Wnt4-Gal4* driving G-TRACE^ts^ stained for DAPI (blue), Fas3 (white), RFP (magenta), and GFP (green). RFP (magenta) marks cells expressing *Wnt4-Gal4*. GFP (green) in addition marks offspring cells of *Wnt4-Gal4* positive cells and can be detected in all follicle cells. Inset shows *Wnt4-Gal4^low^ Fas3^+^* cell at the boundary of *Fas3* expression. C) Quantification of ovarioles with *Wnt4-Gal4* driving G-TRACE^ts^ that do not have GFP-positive cells (unmarked), have only IGS cell clones, transient follicle cell clones, mosaic labeling of the follicle epithelium or a fully marked follicle epithelium. Note that ovarioles at 18°C often contained few GFP^+^ IGS cells, where *Wnt4-Gal4* activity is strongest, but never displayed GFP^+^ follicle cell clones. n=425 ovarioles. D) Heatmap of gene expression in IGS populations, FSCs and pFCs. Some stem cell markers shown in A) show high expression in IGS cells, whereas others are not expressed in IGS cells. See Fig. S5E for corresponding tSNE plot. E) Germarium with *stl-Gal4* driving G-TRACE^ts^ stained for DAPI (blue), Fas3 (white), RFP (magenta), and GFP (green). *stl-Gal4* drives RFP expression sparsely in pFCs in the 2b region, and in differentiated follicle cells. GFP+ clones typically include pFCs in region 2b (inset) but not FSCs or other cells at the 2a/2b border. F) Quantification of ovarioles with *stl-Gal4* driving G-TRACE^ts^ that do not have GFP-positive cells (unmarked), have FSC clones, transient follicle cell clones located in Region 2b, or transient follicle cell clones located posterior to Region 2b. The 18°C control ovarioles frequently contained small GFP^+^ clones of up to 4 cells located posterior to region 3. These clones were usually confined to stalk cells where *stl-Gal4* activity is strongest.

### The Wnt4-Gal4 expression pattern specifically identifies FSCs

The CellFindR results identified *Wnt4* as a marker with strong average expression in the IGS cell clusters and weak average expression in the FSC/pFC cluster (Fig. 1D). Interestingly, the Monocle analysis predicted that, within the FSC/pFC cluster, *Wnt4* is expressed specifically at the beginning of the pseudotime trajectory, and then downregulated at a later stage in the trajectory (Fig. 5A). To test this prediction, we combined *Wnt4-Gal4* with G-TRACE^ts^ and assayed for RFP expression, which marks current driver activity, and GFP expression at 7 dpts. Indeed, we observed strong RFP expression in all IGS cells, as predicted, and significantly lower RFP expression in just 2.08 ± 0.8 cells per germarium (n=79 germaria). Nearly all of the *Wnt4-Gal4^low^* cells (97.7%, n=130 cells) were in a position at the edge of the *Fas3* expression boundary where the FSCs are expected to reside (Fig. 5B). We found that 74.0% of the *Wnt4-Gal4^low^* cells in this position were *Fas3^+^*, which is consistent with previous studies indicating that FSCs reside at the *Fas3* expression boundary and are typically (but not always) *Fas3^+^*.^42, 46^ The possibility that *Wnt4-Gal4^low^* expression is specific for FSCs is particularly interesting because a marker that is capable of distinguishing the FSCs from both IGS cells and pFCs has not been described, and the number and location of the FSCs have been recently debated. Our most recent study directly addressed the controversy of FSC number, and concluded that there are 2-4 FSCs per ovariole,^40^ which aligns well with the observation that only 2 cells per germarium are *Wnt4-Gal4^low^*. To test whether *Wnt4-Gal4* is expressed in FSCs, we looked at the expression of the clonal G-TRACE marker, GFP, in the same ovarioles. Indeed, we observed large GFP^+^ FSC clones that extended through the germarium and across multiple follicles (and thus must have originated from an FSC) in over 97% of the ovarioles at 7 and 14 dpts, including many in which all of the follicle cells in the ovariole were GFP^+^ (Fig. 5B-C). *Wnt4-Gal4* also produced GFP^+^ IGS cell clones, but several lines of evidence argue against the possibility that the GFP^+^ FSC clones originated from the *Wnt4-Gal4^high^* cells. First, *fax-Gal4*, which is expressed sporadically throughout the IGS cell population, including in IGS cells that are adjacent to the *Fas3* border, almost never produced FSC clones (Fig. S4F). Second, *Wnt4-Gal4* is expressed so strongly in IGS cells that it frequently induced IGS cell clones with G-TRACE^ts^ even while the flies are maintained at 18°C (Fig. 5C, “0 dpts” timepoint) but these clones never contributed to the FSC lineage. Lastly, cells with high levels of *Wnt4* expression are transcriptionally distinct from cells with low *Wnt4* expression (compare the first three columns to the fourth column in Fig. 5D) and are sorted into distinct cell clusters by CellFindR (Fig. 4A, Cluster 3.0.1 vs Clusters 3.1.0 and 3.1.1), suggesting that the *Wnt4^high^* IGS cells are not the same cell type as the *Wnt4^low^* cells. Taken together, these results strongly suggest that low *Wnt4-Gal4* expression specifically marks the FSCs.

Our finding that *GstS1* and *Wnt4* are expressed in FSCs as well as IGS populations prompted us to assess the expression of other predicted FSC markers in IGS cells. Interestingly, the majority of genes that Monocle predicts are upregulated in FSCs relative to pFCs (Fig. 5A) are also highly expressed in one or more IGS cell subpopulation (Fig. 5D, S5E). These findings demonstrate that IGS cells and FSCs have significant similarities in expression. However, we also identified 3 genes (*CG31109*, *CG31650*, and *Su(P)*) that are predicted to be highly expressed specifically in the FSCs.

### Monocle identifies novel markers of pFCs

Moving through FSC/pFC pseudotime, after the expression of markers of the FSCs and early pFCs start to decrease, the next stage is distinguished by the upregulation of polar cell markers, such as *upd1* and *neuralized* (*neur*) (Fig. 5A). This is consistent with several previous reports showing that polar cell differentiation is the earliest cell fate decision made by pFCs.^38, 46, 47^ Polar cells induce neighboring pFCs to differentiate into stalk cells and, indeed, the next stage of differentiation in pseudotime was characterized by the upregulation of the stalk cell marker, *CG46339*. We also noticed that *Pdk1*, which we describe above as a marker of anterior and central IGS cells, is upregulated in stalk cells (Fig. 5A, S5D). The final stages of pseudotime are characterized by a series of changes in gene expression, with specific genes peaking in expression at progressively later times. This is consistent with the stepwise progression of pFC differentiation that has been described previously.^8^ One of the earliest markers to peak in expression is *stall* (*stl*) and, indeed, we found that *stl-Gal4* driving G-TRACE^ts^ produced RFP expression starting in Region 2b pFCs (Fig. 5E) but rarely in *Fas3^+^* cells at the Region 2a/2b border, where the FSCs reside. Interestingly, although *stl-Gal4* driving G-TRACE^ts^ frequently produced pFCs clones in Region 2b, it rarely produced FSC clones (Fig. 5E-F), suggesting that pFCs located even just one or two cell diameters downstream from the *Fas3* border do not normally participate in FSC replacement events. The latest marker to peak in expression is *broad* (*br*), and we find that a GFP trap in the Z2 domain (*br*[Z2]*-GFP*) exhibits expression starting in Region 3 pFCs (Fig. 5A, S5F). Notably, *br* has been well-studied at later stages of oogenesis using anti-br antibodies.^48^ However, an anti-BrC antibody that detects the core domain but not the Z1, Z2 and Z3 domains does not detect a signal in the germarium, which may be why *br* expression at this stage was not described earlier. These observations indicate that the wave of expression changes in this portion of the Monocle plot spans the stages of pFC differentiation from Region 2b to Region 3 of the germarium. In addition, our lineage tracing experiment in combination with other studies ^49^ demonstrates that while FSCs can be replaced by pFCs, not all pFCs are fit for competition. This provides evidence for heterogeneity among these transit amplifying cells of the follicle cell lineage.

### Follicle cells exhibit distinct transcriptional profiles corresponding to both stage and position on the follicle

The remaining three Tier 1 clusters (Cluster 0-2) correspond to different stages of main body follicle cells (Fig. 6A-C) and are distinguished from their precursor pFCs (in Cluster 3.0.1) by the lack of *zfh-1* expression (Fig. 6D, S6A). We found that *zfh-1* expression in pFCs diminishes in fully-formed Region 3 cysts (Fig. 4H), indicating that the youngest main body follicle cells in Clusters 0-2 are located in Region 3. Clusters 0, 1 and 2 are broadly distinguished from each other by the enrichment for cells expressing *N-cadherin* (*CadN*) and *Fasciclin 2* (*Fas2*) in Cluster 0, cells expressing *br* in Clusters 1 and 2, and cells expressing *Sox14* in Cluster 2 (Fig. 6B-E, S6B-E). We assayed for expression of these genes by immunofluorescence and found that *CadN* and *Fas2* are expressed in follicle cells in the germarium and early follicles, until Stage 5-6 (Fig. 6H-I), which is approximately when br expression first becomes detectable in differentiated main body follicle cells (Fig. 6J), and Sox14-GFP expression begins at Stage 9 (Fig. S7A). This indicates that Cluster 0 corresponds to early stage main body follicle cells, Cluster 1 corresponds to mid-stage main body follicle cells, and Cluster 2 corresponds to late stage main body follicle cells.

**Figure 6:**
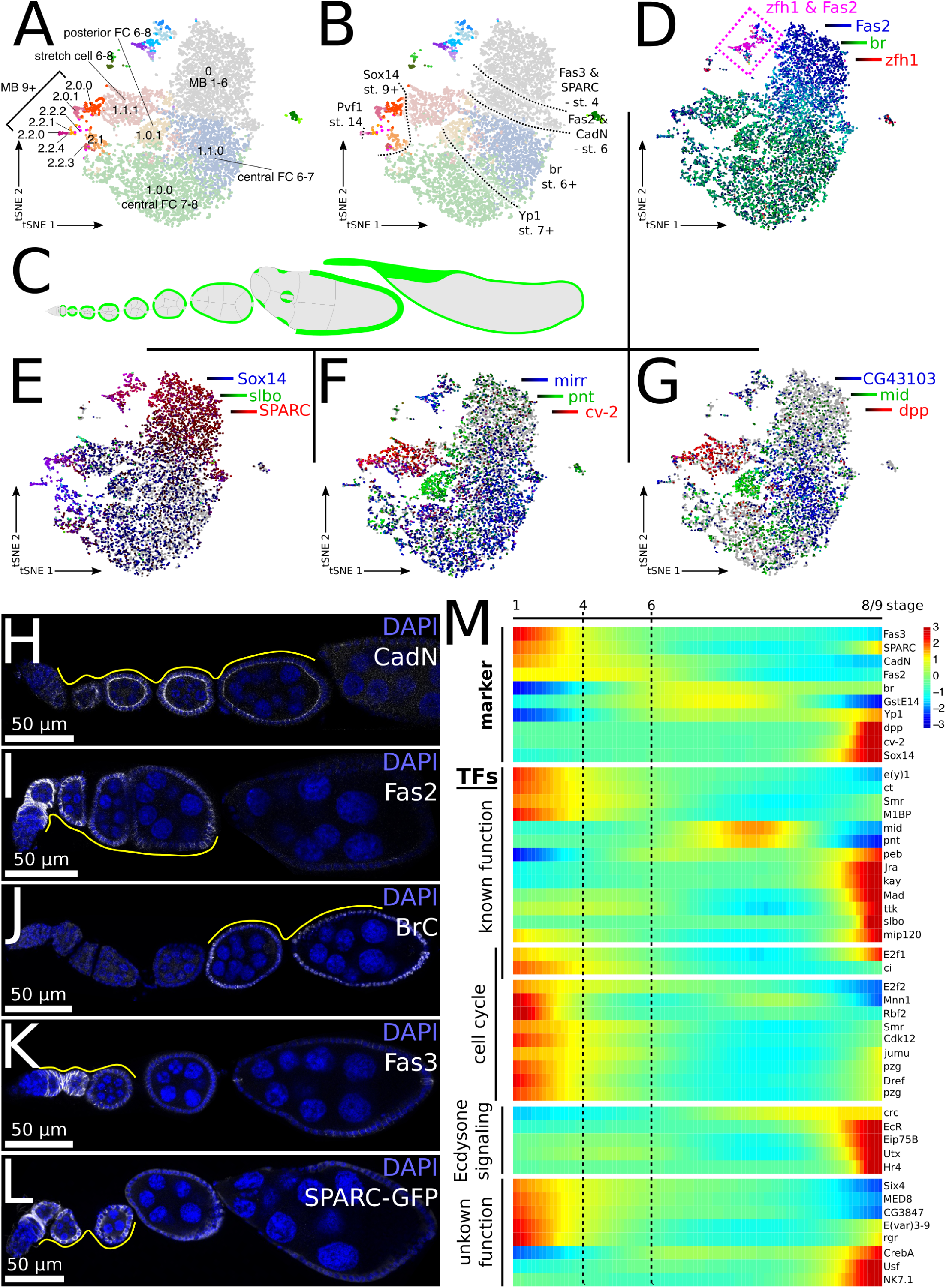
Distinct stages of main body follicle cells. A-B) CellFindR tSNE plots showing main body follicle cell cluster numbers (A) and identities (B). B) Dotted lines mark expression borders for markers expressed in specific stages during main body follicle cell development. Fas3 and SPARC are expressed until Stages 3-4, Fas2 and CadN are expressed until Stages 5-6, br is expressed from Stages 5-6, in Stage 7 central and posterior follicle cells start to express Yp1, Sox14 is expressed from Stage 9 and Pvf1 is detectable in mature Stage 14 main body follicle cells. Note that we approximate stages for several genes due to gradual increase or decrease of expression and potential discrepancy between RNA and reporter line expression. C) Diagram of the ovariole with the main body follicle cells in green. D-G) SCope plots of selected markers of main body follicle cell populations. H-L) Wildtype ovarioles stained for DAPI (blue), and the gene of interest (white). Yellow line outlines positive stages. M) Heatmap of markers and selected transcription factors (TFs) with stage-specific expression in pseudotime of main body follicle cells of Stages 1 - 8/9.

Starting at Stage 6 of oogenesis, follicle cells at the anterior and posterior poles of the follicle acquire distinct identities, and we found that the third tiers of the mid-stage follicle cluster (Cluster 1) correspond to differences in both follicle age and the anterior/posterior position within the follicle. For example, Clusters 1.1.0 and 1.0.0 are distinguished from Cluster 1.0.1 by positional markers, with cells in Clusters 1.1.0 and 1.0.0 expressing *mirror* (*mirr*) (Fig. 6F, S6F), which is a marker of central main body follicle cells in Stages 6-8,^50, 51^ and cells in Cluster 1.0.1 express *midline* (*mid*) and *pointed* (*pnt*), which are markers of posterior follicle cells (Fig. 6I-J, S6G-H, S7B).^52–56^ The two *mirr^+^* clusters (Clusters 1.1.0 and 1.0.0) are distinguished from each other by temporal changes in gene expression, with Cluster 1.0.0 enriched for cells expressing *Yp1*, which is first expressed from Stage 7 (Fig. S6I, S7C). This indicates that the cells in Cluster 1.0.0 are older than the cells in Cluster 1.1.0. Additional genes, such as the expression of *CG43103* in Cluster 1.1.0 but not Cluster 1.0.0 (Fig. 6G, S6J) provide further evidence that these two clusters are distinct populations. The fourth Tier 2 cluster in Cluster 1, Cluster 1.1.1, is enriched for cells expressing *decapentaplegic* (*dpp*) which marks anterior follicle cells, including stretch cells and their precursors (Fig. 6G, S6K, S7D-E)^57^, and *cv-2* (Fig. 6F, S6L, S7F), which is a novel marker of this cell population.

A subset of *dpp* positive cells in the anterior follicle cell cluster (Cluster 1.1.1) expresses *slow border cells* (*slbo*), which is a specific marker of border cells^58^ (Fig. S8A). Border cells are a highly specialized cell type and, indeed, we identified 79 additional genes that are significantly enriched in *slbo^+^* cells (p ≤ 0.05), including *methuselah like-9*, which our analysis predicts is a highly specific marker of border cells (Fig. S8B, D). Border cell differentiation is induced by anterior polar cells, and interestingly, we found that a subset of cells in the polar cell cluster (Cluster 3.2) also express *slbo* (Fig. S8A). In accordance, various predicted border cell specific genes also exhibited expression in a subset of polar cells, including *CG14223* (Fig. S8C-D), which provides additional evidence that the *slbo^+^* polar cells are transcriptionally distinct from other cells in Cluster 3.2, and suggests that they are the anterior polar cells that induce border cell migration.

Cluster 2 is distinguished from the other clusters by the enrichment for cells expressing *Sox14* (Fig. 6E, S6E), which is detectable in follicle cells starting at Stage 9 (Fig. S7A). CellFindR parsed Cluster 2 into a total of 8 clusters (Fig. 1D, S7H). Cluster 2.0.0 and 2.0.1 express the stretch cell marker *dpp* and *cv-2* (Fig. 6F-G, S7I-J). Cluster 2.0.1 is enriched for *Vha16-1* (Fig. S7K,N), which is expressed in stretch cells starting at Stage 10 to induce nurse cell death,^59^ and thus contains older stretch cells than Cluster 2.0.0. Cluster 2.1 is highly enriched for cells that express *Yp1* (Fig. S7L,N), which marks the central and posterior main body follicle cells (Fig. S7C). Monocle analysis of the five clusters in Cluster 2.2 (Cluster 2.2.0 - 2.2.4) predicted a linear lineage relationship, with the youngest cells in pseudotime expressing *Yp1^+^*, and the oldest cells in pseudotime expressing *Pvf1^+^*, which we found is predominantly expressed in the follicle cells of mature Stage 14 egg chambers (Fig. S6M, S7G, M-Q, Table S8). This indicates that these clusters contain follicle cells from follicles that are progressing through the late stages of oogenesis up to Stage 14.

Since follicles follow a developmental trajectory, we applied Monocle to young and mid-stage follicle cells in Cluster 0 and 1 to assay for transcriptional changes that occur during follicle development (Table S9). Monocle placed the cells from early-stage follicles that express high levels of *CadN* and *Fas2*, at one end, and cells from mid-stage follicles that express high levels of *br* at the other end. The cells placed latest in pseudotime started upregulating *Sox14*, in accordance with the expression of *Sox14* in late-stage follicle cells in Cluster 2 (Fig. 6M). In addition, Monocle made predictions about the stage-specific expression of several other genes. For example, Monocle predicted that *Fas3* and *SPARC* expression decrease in early stages of follicle development (Fig. 6M) and, indeed, we found that expression of *Fas3* and *SPARC* both tapered off by Stage 3-4 (Fig. 6F-G), consistent with previous findings.^44, 60^ Interestingly, the *Fas3^+^*, *SPARC*^+^ cells were primarily located near the top of Cluster 0 (Fig. S6N-O), suggesting that the spatial arrangement within this cluster reflects differences in developmental stages.

The arrangement of cells in Cluster 0 and 1 in pseudotime makes it possible to query the dataset for stages of oogenesis that are enriched for specific genes of interest. To test this capability, we searched for stage specific transcription factors and identified 173 genes that fit these criteria (Table S10). Among these we identified several cell cycle regulators that are enriched at the early stages of oogenesis, when follicle cells are mitotic, as well as transcriptional regulators with known functions in oogenesis (Fig. 6M). For example, *E2F1* is a cell cycle regulator that is expressed at early stages to support follicle cell mitosis, then decreases in expression during mid-oogenesis and is re-expressed in late stages to support eggshell chorion gene amplification.^61^ In addition, *ct*, which is downregulated at Stage 6 to allow the mitotic cycle/endocycle switch,^62^ is detectable during early stages only, whereas ecdysone responsive genes are strongly upregulated at Stage 8-9 in accordance with ecdysone signaling steadily increasing from early Stage 9.^63^ Lastly, we identified several transcription factors with unknown functions in oogenesis that will be interesting subjects of follow up studies.

### Posterior IGS cells convert to FSCs in response to physiological stress

The identification of Gal4 lines that are expressed in subsets of IGS cells (Fig. 3) provided us with a unique opportunity to investigate functional differences among cells in the IGS population. As described above, *fax-Gal4* is expressed sporadically throughout the IGS cell population and, in all but one fly (n = 5), did not produce G-TRACE clones in the follicle epithelium (Fig. 3S, Fig. S4F). Our finding that the only four ovarioles with clones in the follicle epithelium were all isolated from the same fly prompted us to consider whether environmental conditions such as nutrient availability could affect the pattern of clone formation. Indeed, we found that exposure to 24 hours of starvation in the middle of a 14-day period with *fax-Gal4* driving G-TRACE produced FSC clones in 80% of flies examined (n = 5). When we quantified all ovarioles from each fly in aggregate, we found that 6.2% of ovarioles contained FSC clones, and an additional 3.3% of ovarioles contained transient follicle cell clones (n=211). In contrast, we did not find any follicle cell clones when flies were kept on rich diet throughout the 14 days (n = 5 flies, n=170 ovarioles) (Fig. 7A-B, E). Starvation conditions did not expand the expression of RFP into the *Fas3^+^* region (Fig. S9A-C), indicating that the emergence of clones is not due to the expression of *fax-Gal4* in follicle cells. Consistent with this, we confirmed previous reports ^29^ that starvation causes a decrease in *fax-GFP* expression in IGS cells (Fig. S9D-F).

**Figure 7:**
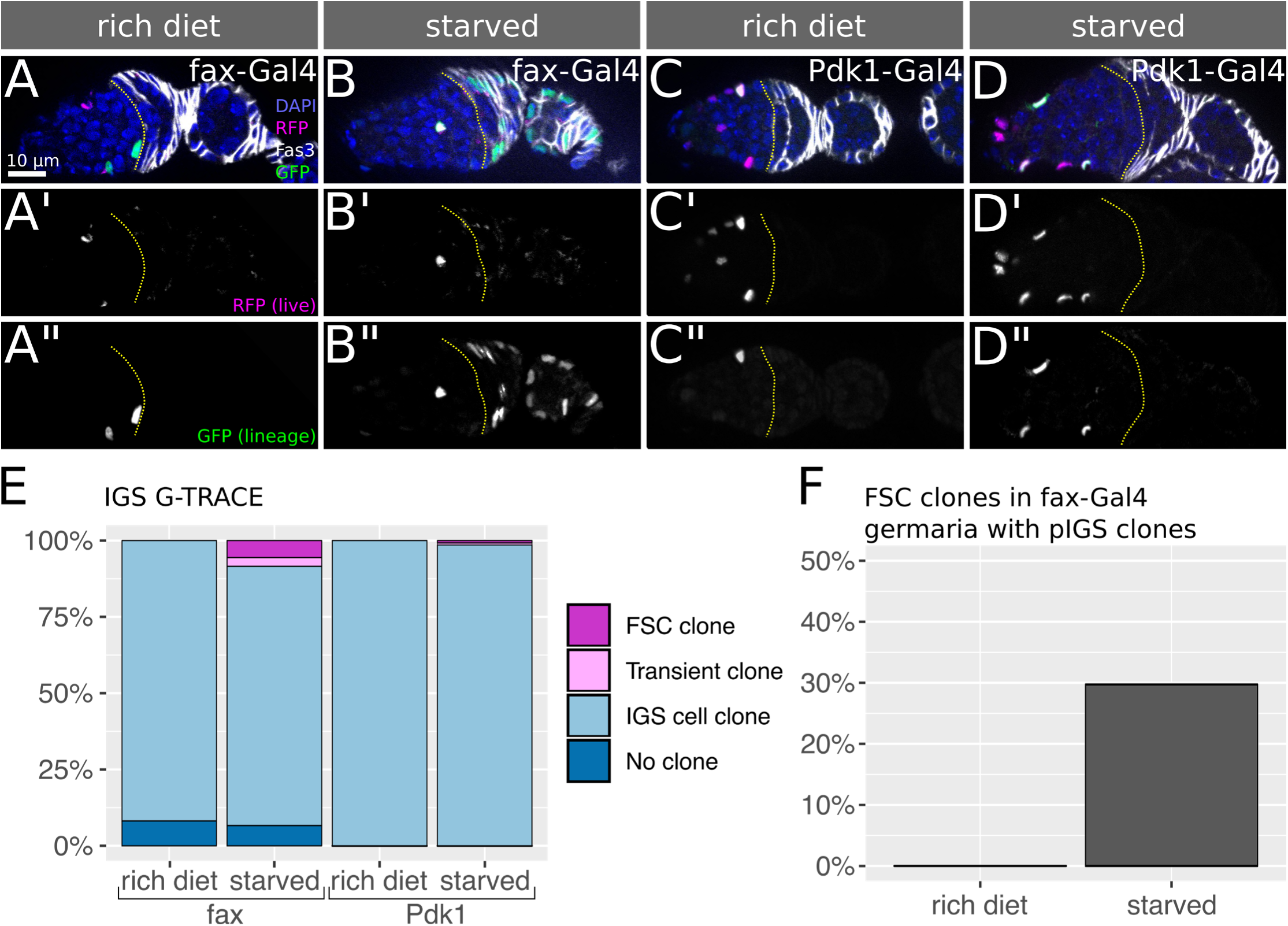
Posterior IGS cells convert to FSCs in response to nutrient deprivation. A-D) Germaria from flies with *fax-Gal4* (A-B) or *Pdk1-Gal4* (C-D) driving G-TRACE stained for DAPI (blue), Fas3 (white), RFP (magenta), and GFP (green). Yellow line outlines the 2a/2b border. *fax-Gal4* and *Pdk1-Gal4* do not produce GFP^+^ FSC clones on a rich diet (A,C), whereas, following exposure to 24h of starvation, *fax-Gal4*-expressing cells produce GFP^+^ FSC clones (B) but *Pdk1-Gal4*-expressing cells almost never do (D) (we observed 1 ovariole with a transient clone and 1 ovariole with an FSC clone out of 138 ovarioles total). E) Quantification of GFP^+^ clone types in flies with *fax-Gal4* or *Pdk1-Gal4* driving G-TRACE^ts^ exposed to a rich diet or starvation. n=170, 211, 134, and 138 ovarioles for *fax-Gal4* rich diet, *fax-Gal4* starved, *Pdk1-Gal4* rich diet, and *Pdk1-Gal4* starved, respectively. F) Quantification of FSC clones in *fax-Gal4* > G-TRACE^ts^ germaria that contained at least one GFP positive IGS cell adjacent to the Fas3 border. n=20 in rich diet, n=37 under starvation.

With *Pdk1-Gal4*, which is not expressed in pIGS cells (Fig. 3J,L,T-U), we found that flies exposed to identical starvation conditions did not produce FSC or follicle cell G-TRACE clones in 4 out of 5 flies examined. In the one fly with clones, one ovariole had an FSC clone and one had a transient clone (0.7% of total, n=138 ovarioles) (Fig, 7C-E). *Pdk1-Gal4* is expressed in many more IGS cells per germarium than *fax-Gal4* (Fig. 3S-T) but, in contrast to *fax-Gal4*, only rarely produces GFP^+^ pIGS clones (Fig. 3S-U, 7A,C). Together, this suggests that pIGS cells located adjacent to the Fas3 boundary can convert to follicle stem cells under starvation conditions while aIGS and cIGS cells located closer to the anterior cannot. We reasoned that only germaria with a GFP^+^ pIGS could display FSC clones after starvation. Therefore, to estimate the rate of IGS to FSC conversion under starvation more precisely, we analyzed the occurrence of FSC clones in *fax-Gal4* containing at least one GFP^+^ pIGS cell. We found that, whereas none of the germaria with GFP^+^ pIGS cells contained FSC clones at 14 dpts (n=20) in flies that were maintained on a rich diet, 29.7% (n=37) of these germaria from flies exposed to 24 hours of starvation contained FSC clones (Fig. 7F). Taken together, these observations strongly suggest that pIGS cells, but not aIGS and cIGS, cells are able to convert to FSCs in response to starvation.

## Discussion

In summary, we have generated a detailed atlas of the cells in the Drosophila ovary. This atlas consists of 26 groups of cells that each correspond to a distinct population in the ovary (Fig. S10A-B, Table S11). Through experimental validation and referencing well-characterized markers in the literature, we determined the identity of each group, and found that all of the major cell types in the ovariole are represented. We further identified several transcriptionally distinct subpopulations within these major cell types, such as the anterior, central, and posterior IGS cell populations. We also identified both the GSCs and the FSCs in our dataset, which revealed several genes that are predicted to be specific for each of these stem cell populations. In addition, we identified several Gal4 lines, including *Pdk1-Gal4*, *fax-Gal4*, and *stl-Gal4*, with unique expression patterns that make it possible for the first time to target transgene expression to the corresponding subset of cells (such as *Pdk1-Gal4* for aIGS and cIGS cells). Lastly, although we have primarily focused on the most uniquely expressed genes for each cluster in this study, the broad transcriptional profile of each cluster is a rich dataset that can be mined to identify populations of cells that are relevant for a topic of interest. For example, we compared the gene expression profile of each cluster to a list of human disease genes that are well-suited for analysis in Drosophila.^64^ We found that germ cells are enriched for cells expressing major drivers of cancer, and IGS and follicle cells are enriched for genes involved in cardiac dysfunction (Fig. S10C), suggesting that these cell types may be good starting points for studies into the genetic interactions that underlie these human diseases.

This study also demonstrates the utility of using CellFindR^13^ in combination with Monocle^23^ to identify unique populations of cells within a large dataset. Because CellFindR produces clusters in a structured, iterative fashion, we were able to construct a hierarchical tree that corresponds to a transcriptome relationship between clusters, and this outperformed other hierarchical clustering methods (Fig. S1N). The tree built by CellFindR aligns well with expectations and provides some interesting new insights. For example, we expected that germ cells would cluster away from somatic cells in Tier 1 because these populations are substantially different from each other, arising at different times during development and from completely different lineages. However, it was surprising that the FSC, pFCs, polar cells and stalk cells clustered more closely to IGS cells than to the follicle cells of budded follicles. This suggests that many cell types in the germarium, which are often studied separately, have biologically relevant similarities. In the budded follicles, we found that the Tier 2 and 3 distinctions are driven by both stage of oogenesis and position in the follicle, thus describing two separate axes of differentiation in these populations. Overall, the distinctions between clusters at every tier correspond well with the biology of the tissue, demonstrating the accuracy of the procedure.

Using Monocle, we were able to identify more subtle distinctions within the clusters. Interestingly, we found that Monocle was not only able to build informative trajectories based pseudotime, as it is typically used for, but also based on changes in pseudodistance across a population of cells (the IGS cells) with a graded pattern of gene expression. The pseudotime trajectories made it possible to distinguish the GSCs and FSCs from their daughter cells as well as distinct stages of differentiation in the main body cell population, whereas the pseudodistance trajectory identified at least three distinct populations of IGS cells. However, since Monocle identified five distinct states of IGS cells, it is possible that there is even more diversity in the IGS cell population than we were able to resolve in this study.

Our analysis of the FSC transcriptome led to the identification of *Wnt4-Gal4*, which is the first marker that uniquely identifies the FSCs. FSCs are known to reside at the boundary of *Fas3* expression, though the FSCs themselves appear to exhibit variable levels of *Fas3* expression.^46^ In addition, this criterion alone is not sufficient because there are 8-10 cells at the *Fas3* expression boundary but there are only 2-4 FSCs per germarium.^40^ Moreover, *Fas3* is also expressed in pFCs, so it is not useful for testing whether a particular experimental manipulation interferes with the transition from the FSC to pFC fate. Likewise, low levels of *Castor* and *Eyes absent* expression have been reported to mark cells at the boundary of *Fas3* expression,^38, 65, 66^ but have not been shown to distinguish between the 8-10 cells at this boundary. In addition, like *Fas3*, both genes are also expressed in pFCs, and the differences in expression levels between the FSCs and pFCs are subtle. Nonetheless, these three markers are useful for distinguishing follicle cells from IGS cells and are commonly used for this purpose. Transcriptional reporters of Wnt pathway activity and phosphorylated ERK are strongly upregulated in FSCs compared to pFCs^30, 33, 38, 67, 68^ but these markers are equally strong in the neighboring IGS cells, and thus are also not specific for the FSCs. In contrast, we consistently found that low levels of *Wnt4-Gal4* expression marked just 2-3 cells at the boundary of *Fas3* expression, which fits well with the number of FSCs we would expect in this position, and we provide functional evidence with lineage tracing that *Wnt4-Gal4* is expressed in FSCs. Although *Wnt4-Gal4* is also strongly expressed in IGS cells, we describe bioinformatic distinctions between *Wnt4-Gal4^high^* and *Wnt4-Gal4^low^* cells. Further, additional lineage tracing data using the IGS specific driver *fax-Gal4* suggest that the *Wnt4-Gal4^high^* cells are a distinct population of cells that do not contribute to the FSC lineage under standard laboratory conditions. Therefore, our data strongly argue that *Wnt4-Gal4^low^* specifically marks the FSCs.

While FSCs express specific markers, pFCs are characterized by a stepwise transcriptional program towards their differentiated fates. This insight into the biology of a transiently amplifying cell population opens new lines of inquiry such as the robustness and responsiveness of this transcriptional program to cellular manipulations. Recent studies suggest that stem cell replacement by the transit amplifying daughter cells can be a driving force of cancer development.^69–71^ Our finding that pFCs lose the ability to compete for the stem cell position after moving just 1-2 cell diameters away from the stem cell niche supports the idea of a “point of no return”, and it will be interesting to investigate the underlying biology of this transition in future studies.

Our use of G-TRACE to assess the lineage potential of somatic cells in the germarium led to the surprising finding that pIGS cells can convert to FSCs under starvation conditions. Recent studies have described other forms of cellular plasticity in stem cell based tissues, suggesting that this may be a well-conserved aspect of tissue homeostasis. For example, in the mammalian intestine, stem cell daughters that have begun to differentiate are capable of reverting back to the stem cell state in response to severe tissue damage or stem cell ablation.^72–75^ Likewise, the cyst stem cells in the *Drosophila* testis produce post-mitotic niche cells that are capable of converting back to cyst stem cells upon RNAi knockdown of the transcription factor, *escargot*.^76, 77^ Our findings reveal a new form of cellular plasticity in which pIGS cells, which are not part of the FSC lineage during normal homeostasis,^7, 40^ can convert to FSCs in response to starvation. This builds on these previous observations by demonstrating that new stem cells can derive from a closely related but independent lineage, and demonstrates that the cell fate conversion can be induced as part of a natural response to a physiological stress. The Gal4 lines that are typically used to drive expression in IGS cells, such as *13C06-Gal4* and *c587-Gal4*, are also expressed in FSCs, so this discovery was not possible without the availability of a driver such as *fax-Gal4*. The frequency of FSC and transient clones produced by *fax-Gal4* driving G-TRACE under starvation conditions is 29.7% in germaria with pIGS clones. However, this is likely an underestimate of the rate at which pIGS cells can convert to FSCs because *fax-Gal4* is a relatively weak driver and usually does not mark all pIGS cells. We demonstrated recently that IGS cells provide a niche-like function by delivering a juxtacrine wingless signal to the FSCs.^33^ It is unclear whether the IGS cells that provide this signal are the same cells that convert to FSCs in response to starvation but, taken together, these observations suggest the interesting possibility that niche cells may be able to convert to stem cells in response to stress.

Overall, this study provides a new resource that will be valuable for a wide range of studies that use the *Drosophila* ovary as an experimental model. Additional scRNA-Seq datasets provided by other studies will further increase the accuracy and resolution of the ovary cell atlas, and it will be important to follow up on the predictions of the atlas with detailed studies that focus on specific populations of cells. Collectively, these efforts will help drive discovery forward by providing a deeper understanding of the cellular composition of the *Drosophila* ovary.

## Supplementary tables

**Table S1:** Markers identified by CellFindR with ≥10 genes

**Table S2:** Markers of subclusters of cluster 1 identified by CellFindR with 5 ≥ genes

**Table S3:** Markers of clusters in replicate sample identified by Seurat

**Table S4:** Markers of germ cells identified by Monocle

**Table S5:** Markers of young germ cells and oocytes identified by Monocle

**Table S6:** Markers of IGS populations identified by Monocle

**Table S7:** Markers of FSCs and pFCs identified by Monocle

**Table S8:** Markers of young and middle-staged main body follicle cells identified by Monocle

**Table S9:** Transcription factors with stage-specific expression in main body follicle cells

**Table S10:** Markers of late-stage follicle cells in Cluster 2.2 identified by Monocle

**Table S11:** Markers of all distinct cell types identified by Seurat

## Material and Methods

### Single-cell sequencing of the Drosophila ovary

Newly hatched Canton-S flies were reared on standard lab conditions and fed wed yeast for three consecutive days. 60 females were dissected within 45 min in ice cold Schneider’s Insect Medium with 10 % FBS and 167 mg/ml insulin on an ice pack. We enriched for the younger, non-vitellogenic stages of the ovary using micro-scissors. Tissue was transferred to an eppendorf tube containing ice cold Cell Dissociation Buffer (Thermo Fisher Scientific #13151014) and rinsed once with the buffer. Dissociation was performed at RT in Cell Dissociation Buffer with 4 mg/ml elastase (Worthington Biochemical LS002292) and 2.5 mg/ml collagenase (Invitrogen # 17018-029) with nutation and regular pipetting with a P200 to aid tissue dissociation. After 20 min the solution was passed through a 50 µm filter (Partec #04-0042-2317) and the solution incubated for additional 10 min before passage through a 30 µm filter (Miltenyi Biotec #130-041-407). Enzymes were quenched by adding 500 µl of dissection solution and cells were centrifuged for 5 min at 4°C and 3500 rcf. Cells were washed in dissection solution and centrifuged again before being resuspended in ice cold 200 µl PBS with 0.04 % ultrapure BSA (Thermo Fisher Scientific AM2616). Dissociation was verified and cells were counted using a cell counting chamber and the solution adjusted to 1000 cells per µl before subjection to single-cell RNA-sequencing using the Chromium Single Cell 3’ Reagent Version 2 Kit (10x Genomics). For the bigger dataset 27000 cells were loaded into one well of the 10X chip. Sequencing was performed on Illumina HiSeq 2500 according to the 10X Genomics V2 manual. For the second, smaller replicate we dissected 200 flies with the genotype *109-30-Gal4/+; 13CO6-GFP/UAS-CD8::GFP* and treated them similarly to receive a single-cell solution which was subjected to MACS as described before^47^ and then subjected to single-cell sequencing using the 10X platform. For this dataset we loaded 5,000 cells.

### Bioinformatic analysis

Reads were aligned to the Drosophila reference genome (dmel_r6.19) using STAR and resulting bam files were processed with the Cell Ranger pipeline v2.1.1 (10K dataset) or v2.0.0 (replicate dataset). For the bigger dataset of initially 10964 cells, 1071 doublets were removed based on number of genes (keeping cells with 1000 - 4000 genes), UMI counts (keeping cells with ≥ 5000 and ≤ 30000 UMI reads) or expression of known mutually exclusive genes using Seurat v3.0.2. We estimate approximately 6700 cells per ovariole, thus this dataset achieves 1.5x coverage. Clustering was performed with CellFindR using standard settings with a quality measure of ≥ 10 genes. For R scripts and complete details about CellFindR, see Yu, et al., 2019. With these parameters, Cluster 1 was not sub-clustered in Tier 2, yet we could clearly identify subpopulations based on previously described markers. Therefore, we repeated CellFindR on Cluster 1 using a quality measure of ≥ 5 genes. Subsequent analysis was performed with Seurat v3.0.2. For a second replicate of 550 cells CellFindR was performed with ≥ 10 genes to identify cluster with high cell number. Cluster which were not identified by CellFindR due to low cell number, were assigned by validated marker expression. To compare CellFindR and Seurat on our 10964 cell dataset, we performed Seurat v3.0.2 with identical filtering, 2000 variable features identified by vst-method and UMAP dimension reduction with 20 dimensions. Pseudotime analysis was performed with Monocle v2.8.0. Multicolor tSNE plots were visualized using SCope.78 Scales in dotplots, expression plots and Monocle plots correlate with the fold-change in transcript levels of the indicated gene.

### Fly husbandry

Flies were reared under standard lab conditions at 25°C and fed wet yeast for at least three consecutive days prior to dissections. For G-TRACE experiments, Gal4-drivers were combined with tub-Gal80ts and bred at 18°C to repress G-TRACE activity during developments. For 18°C controls flies were kept at the restrictive temperature fed wed yeast for at least 3 consecutive days prior to dissection. For 7d and 14d time points adult flies were shifted to 29°C and fed wet yeast daily until dissection. Starvation experiments were conducted at 29°C. Flies were kept for 7d and fed wet yeast daily to allow induction of G-TRACE, starved for 24h in an empty vial with a wet kimwipe, and shifted back to rich diet until dissection. For intensity measurements of fax-GFP, control flies were fed wet yeast for three consecutive days, while starved flies were shifted to an empty vial with a wet kimwipe on day 2 and dissected after 24h starvation.

### Fly stocks

The following fly stocks were used in this study: BDSC stocks: Canton-S (64349), Orc1-GFP (52168), CG44325-GFP (53795), santa-maria-Gal4 (24521), GstS1-lacZ (11036), fax-Gal4 (77520), Pdk1-Gal4 (76682), CG46339-Gal4 (77710), Jupiter-GFP (6825), Wnt4-Gal4 (67449), stl-Gal4 (77732), SPARC-GFP (5611), br^[Z2]^-GFP (38630), Sox14-GFP (55842), Pvf1-lacZ (12286), pnt-GFP (42680), dpp-lacZ (12379), cv-2-lacZ (6342), fax-GFP (50870), G-TRACE: UAS-RedStinger, UAS-Flp, Ubi-(FRT.STOP)-Stinger (28281), tub-Gal80ts (7108), 109-30-Gal4 (7023), UAS-CD8::RFP (27399). 13CO6-GFP (generated from BDSC stock 47860)^30^ VDRC: Yp1-GFP (318746). hh-lacZ and ptc-pelican (kind gifts from Tom Kornberg), upd-Gal4 (kind gift from Denise Montell).

### Immunofluorescence staining and imaging

Flies were dissected in PBS at RT and ovaries were fixed for 15 min at RT with 4% PFA. Ovaries were washed with PBS twice and blocked for 30 min at RT with blocking solution (PBS with 0.2% Triton X-100 and 0.5% BSA). Primary antibody incubation was performed overnight at 4°C in blocking solution. On the following day ovaries were washed three times for 10 min with blocking solution and incubated with secondary antibodies diluted in blocking solution for 4h. After washing with PBS ovaries were mounted in DAPI Fluoromount-G (Thermo Fisher Scientific, OB010020) and imaged using a Zeiss M2 Axioimager with Apotome unit or Nikon C1si Spectral Confocal microscope. Images were analysed with FIJI.^80^ The following antibodies were used in this study:

DSHB: mouse anti-Fas3 (7G10, 1:100), rat anti-CadN (DN-EX#8-s, 1:10), mouse anti-Fas2 (1D4, 1:100), mouse anti-BrC (25E9.D7, 1:50), mouse anti-gro (anti-Gro, 1:1000), mouse anti-en (4D9, 1:25). rabbit anti-GFP (Cell Signaling #2956, 1:1000), guinea pig anti-GFP (Torrey Pines Biolabs Inc #116-67,1:1000), mouse anti-beta Galactosidase (Promega Z378A, 1:100), chicken anti-beta Galactosidase (abcam ab9361, 1:100), rat anti-RFP (ChromoTek 5F8, 1:1000), rabbit anti-vas (Santa Cruz Biotechnology sc-30210, 1:1000), guinea pig anti-tj (kind gift from Dorothea Godt, 1:5000), guinea pig anti-zfh1 (kind gift from Dorothea Godt, 1:500), anti-chicken 555 (Sigma-Aldrich, SAB4600063). Additional secondary antibodies were purchased from Thermo Fisher Scientific and used at 1:1000: goat anti-rabbit 488 (A-11008, goat anti-rat 555 (A-21434), goat anti-mouse 647 (A-21236), goat anti rabbit 555 (A-21428), goat anti-guinea pig 488 (A-11073), goat anti-guinea pig 555, goat anti-mouse 488 (A-11029), goat anti-mouse 555 (A-21424), goat anti-rat 555 (A-21434).

Images were acquired with either a Zeiss M2 Axioimager with Apotome unit or a Nikon C1si Spectral Confocal microscope. Image processing and analysis was performed with FIJI.^79^

## Acknowledgements

We thank Tom Kornberg and Denise Montell for sharing fly lines and Dorothea Godt for antibodies. We are grateful to Marco Conti, Sumitra Tatapudy and Nathaniel P Meyer for critical comments on the manuscript. We also thank the Bloomington Stock Center, the Vienna Drosophila Resource Center and the Developmental Studies Hybridoma Bank for many stocks and resources used in this paper. K.R. is supported by the Deutsche Forschungsgemeinschaft (DFG, project number 419293565), and T.N. is supported by a grant from the National Institutes of Health, GM097158.

---

**Figure S1:**
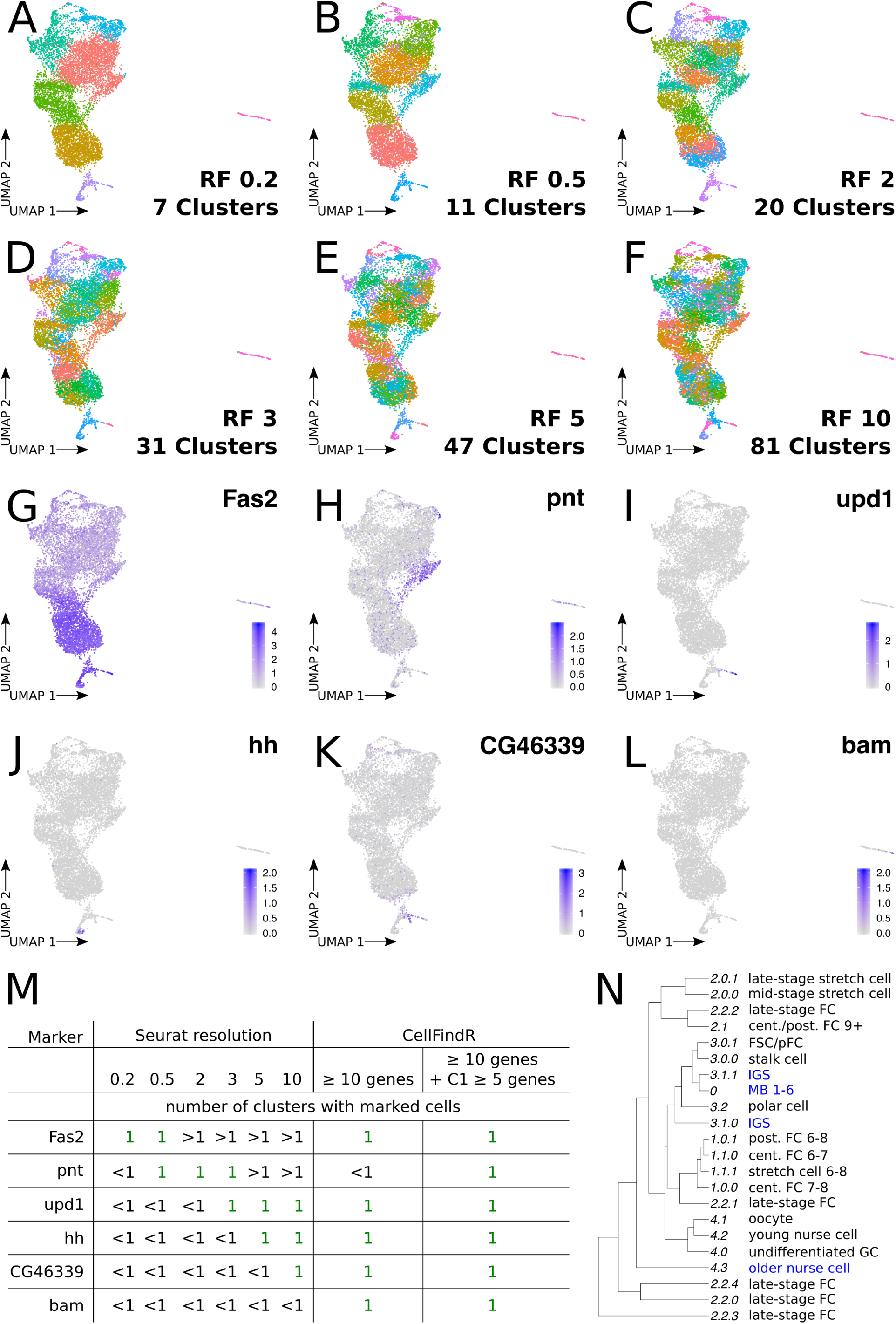
Comparison of Seurat and CellFindR. A-F) UMAP - clustering of 9623 cells isolated from Drosophila ovaries with Seurat v3.0.2 with different resolution factors (RF) without subsequent merging of clusters. Number of resulting clusters is indicated. G-L) Expression of confirmed markers, indicated on panels, of specific cell populations identified by CellFindR on Seurat v3.0.2 UMAP-plot. M) Table comparing the number of clusters for validated markers between Seurat v3.0.2 clustering with different resolution factors to both the initial CellFindR object produced by setting the number of different genes to ≥ 10 genes and to our final clustering where Cluster 1 was sub-clustered with a setting of ≥ 5 genes. Note that even a setting that produced 81 clusters in Seurat was unable to identify the bam+ cluster. N) Cluster tree build with Seurat v3.0.2 of CellFindR clustering as shown in Fig. 1E. Blue font indicates cell types with positions on the tree that do not fit expectations.

**Figure S2:**
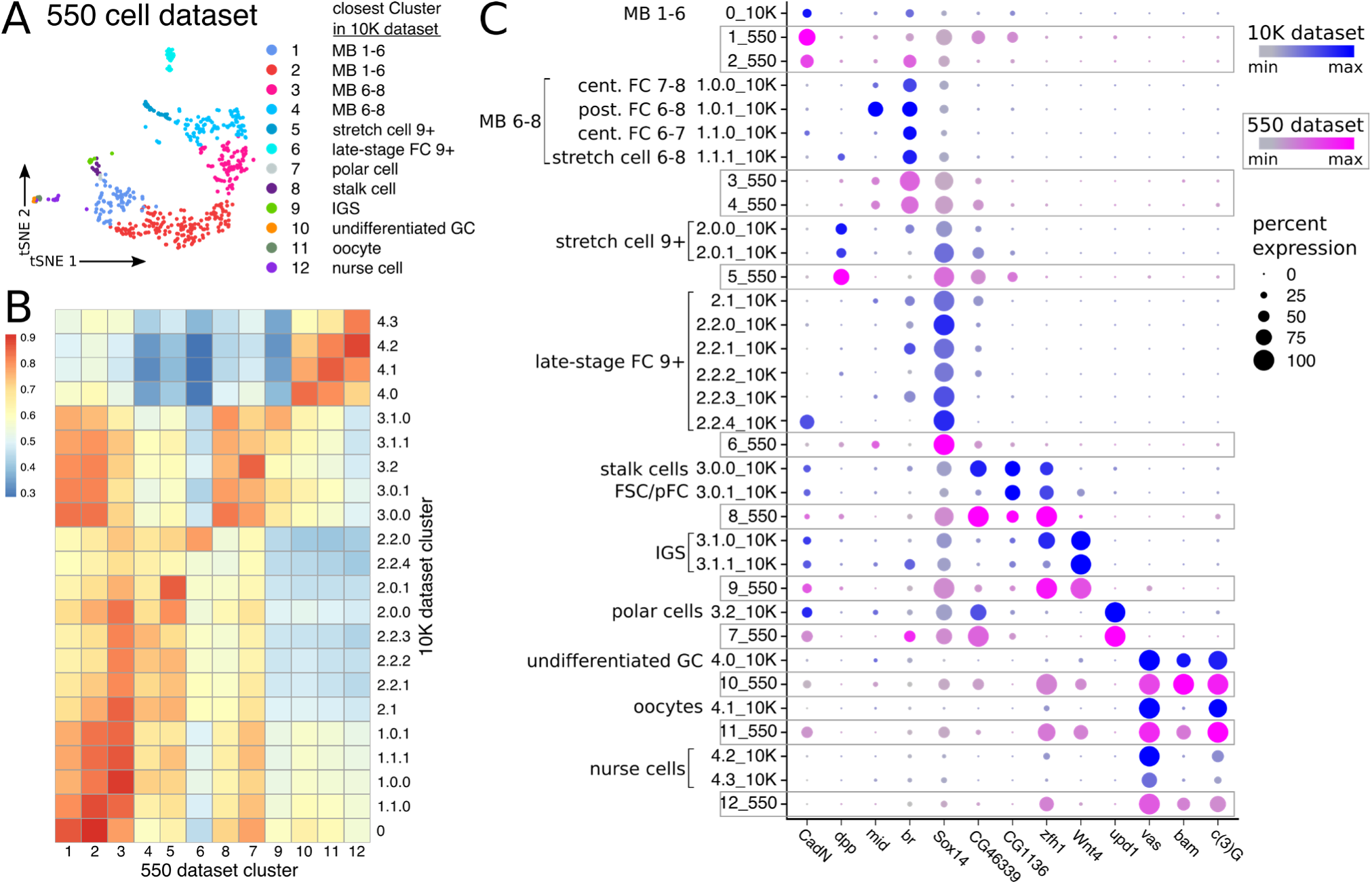
Sample validation. A) tSNE plot of 550 cells isolated from adult Drosophila ovary. B-C) Spearman correlation heatmap of shared variable genes (B) and dotplot of selected markers (C) in two replicates with 9623 cells (10K) and 550 cells (550) isolated from Drosophila ovaries. The two datasets are strongly correlated even though the two datasets were generated using different experimental procedures and were analyzed with slightly different procedures (see Materials and Methods). Note that while the 10K dataset achieved 1.5x coverage, the 550 cell dataset does not contain all cell types, which may explain the discrepancy in cluster numbers.

**Figure S3:**
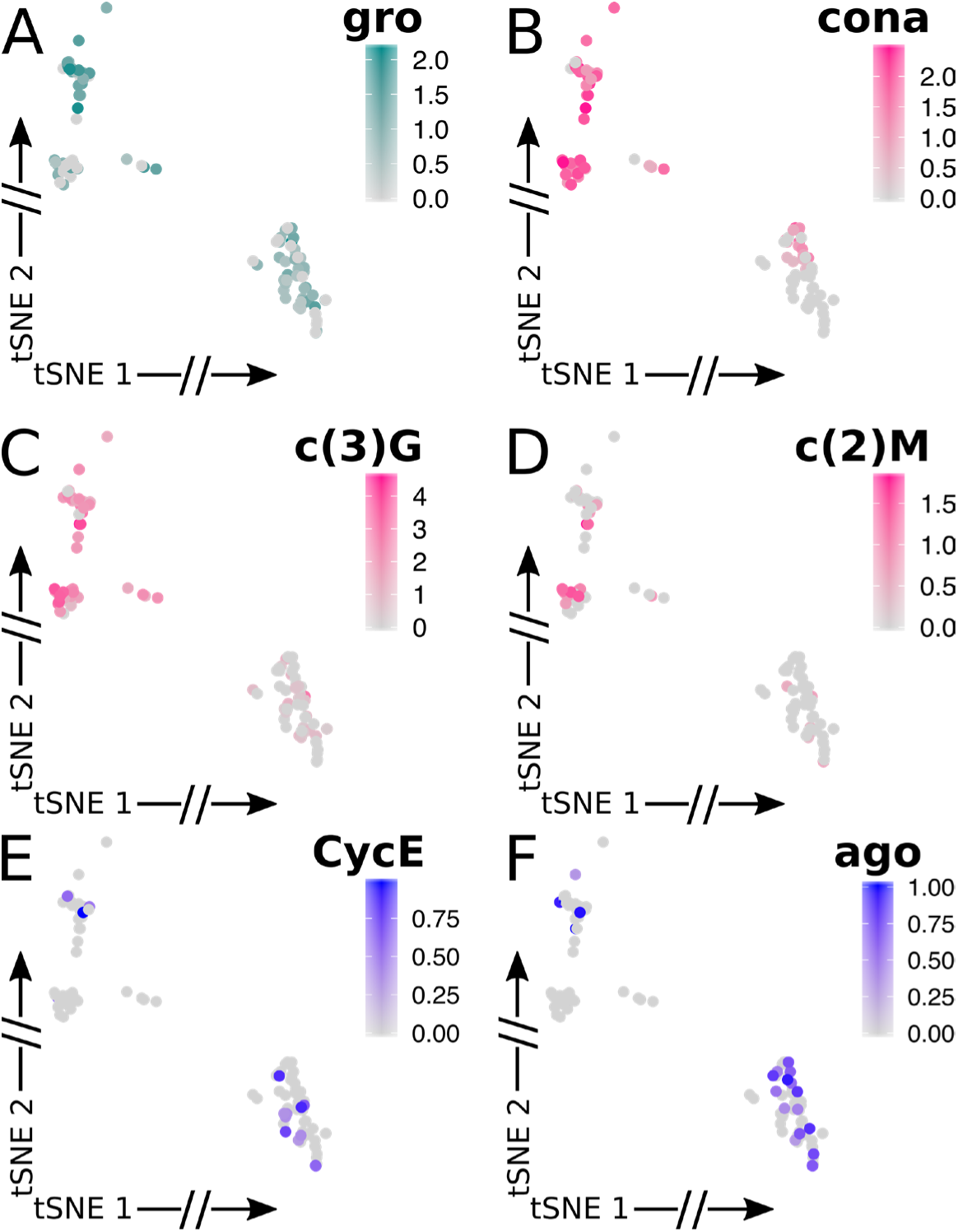
tSNE plots for germ cell markers. A-F) tSNE plots of the germ cell cluster (Cluster 4) showing the expression patterns of the indicated genes. A) Cluster 4.0, is enriched for cells expressing gro. B-D) Clusters 4.0 and 4.1 are enriched for meiotic genes. E-F) Clusters 4.2 and 4.3 are enriched for genes involved in endoreduplication.

**Figure S4:**
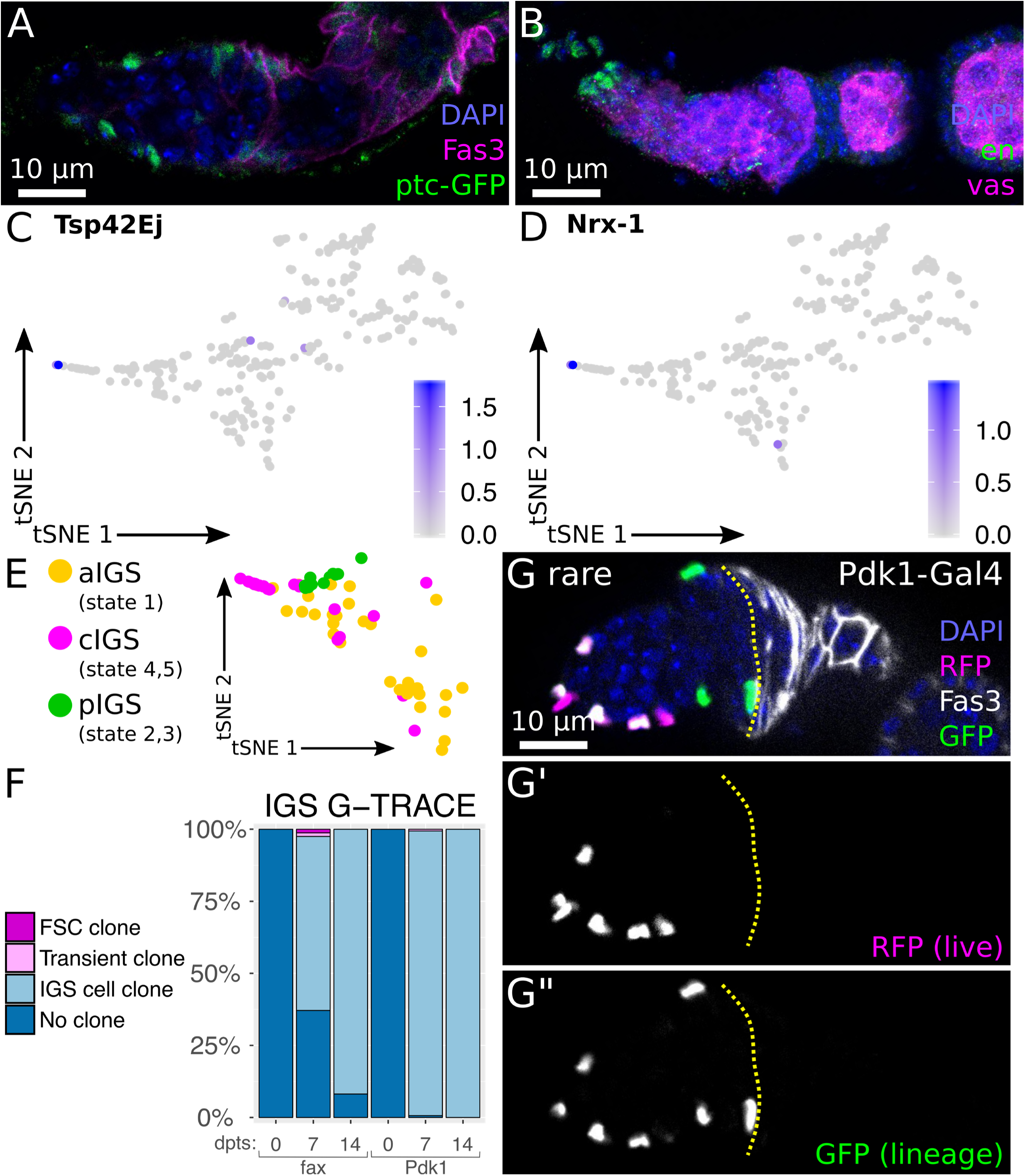
Apical cells and IGS cell lineage tracing. A-B) Immunofluorescence staining of the early stages of the ovariole. DAPI (blue) marks nuclei. A) ptc-GFP (green) is highly expressed in IGS cells. Fas3 (magenta) marks follicle cells. B) Maximum intensity projection. en (green) is expressed in terminal filament and cap cells. vas (magenta) marks germ cells. C-D) Expression of potential terminal filament and cap cell markers. E) tSNE plot showing IGS subpopulations identified by Monocle. F) Quantification of the percentage of ovarioles without GFP positive cells (unmarked), only IGS GFP positive cells (IGS positive), and IGS cells positive with transient follicle cell clones (transient FC) or FSC clones (FSC clone). n = 167,167, and 170 for *fax-Gal4* at 0, 7, and 14 dpts, respectively and 113, 159, 134 for *Pdk1-Gal4* at 0, 7, and 14 dpts, respectively. Note that the rare ovarioles with follicle cell clones derived from a single fly. G) *Pdk1-Gal4* driving G-TRACE. Fas3 (white) marks follicle cells. Yellow line marks 2a/2b border. RFP (magenta) marks *Pdk1-Gal4* positive aIGS and cIGS. GFP (green) is also expressed in offspring cells of formerly *Pdk1-Gal4* positive cells and in rare cases marks cells of all IGS populations.

**Figure S5:**
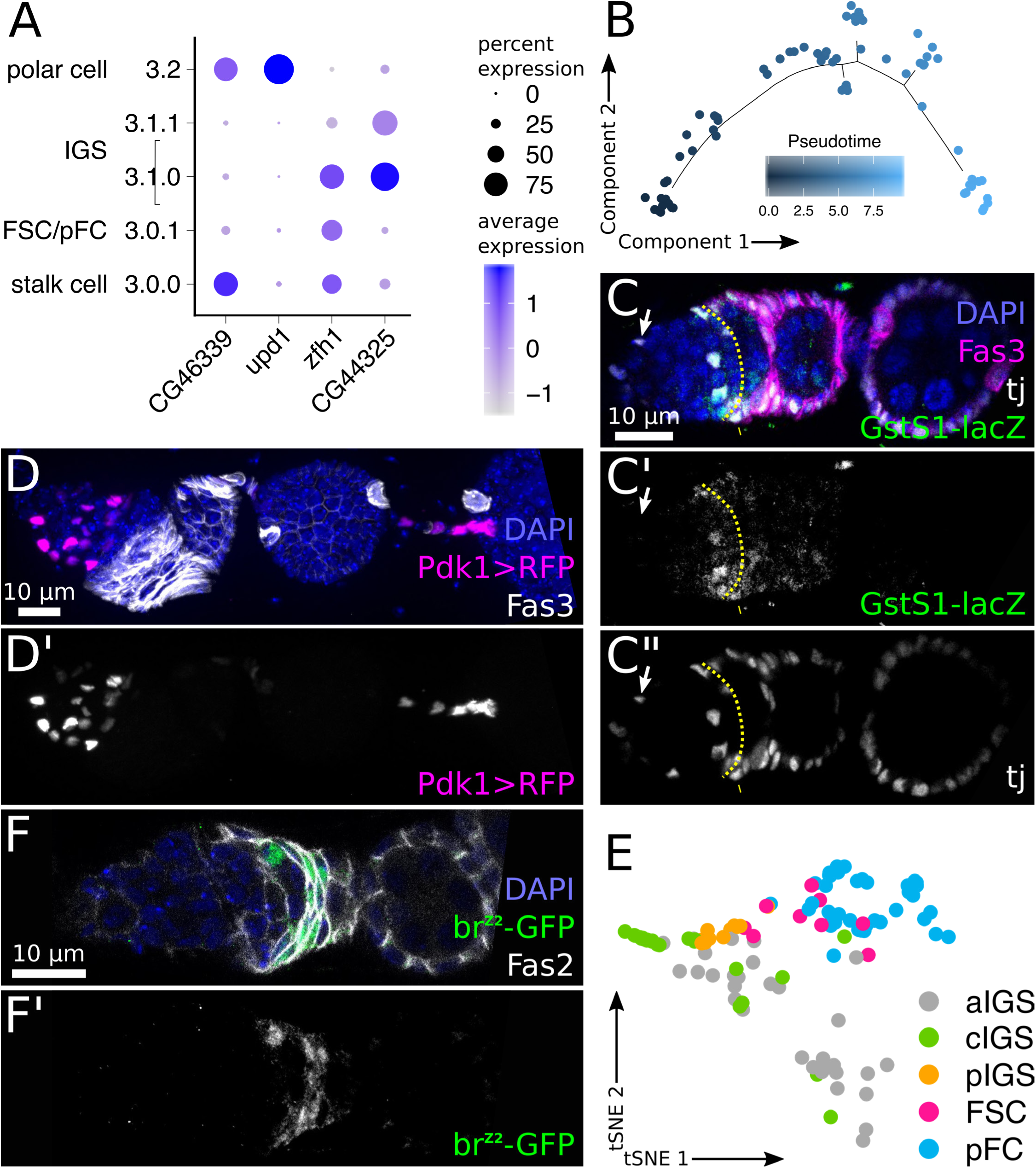
Markers of early stages in the FSC lineage. A) Expression of selected markers in Cluster 3. B) Monocle plot of FSCs and pFCs in Cluster 3.0.1 colored by pseudotime. Compare heatmap in Fig. 5A. C-D) Immunofluorescence staining of young stages of the ovariole. DAPI (blue) marks nuclei. C) Fas3 (magenta) marks follicle cells. The 2a/2b border is marked with a yellow line. tj (white) is expressed in somatic cells. GstS1-lacZ (green) is absent from anterior IGS cells (arrow) and expressed in IGS closer to the 2a/2b border and detectable in a small number of cells positive for Fas3 posterior to the 2a/2b border (arrowhead). D) Pdk1>RFP can be detected in aIGS and cIGS and is expressed in mature stalk cells. E) tSNE plot with aIGS, cIGS, pIGS, FSCs and pFCs. These cell populations were used for the heatmap shown in Figure 5D. F) Immunofluorescence staining. br^[Z2]^-GFP (green) is expressed in pFCs in region 2b. Fas2 (white) is highly expressed in follicle cells.

**Figure S6:**
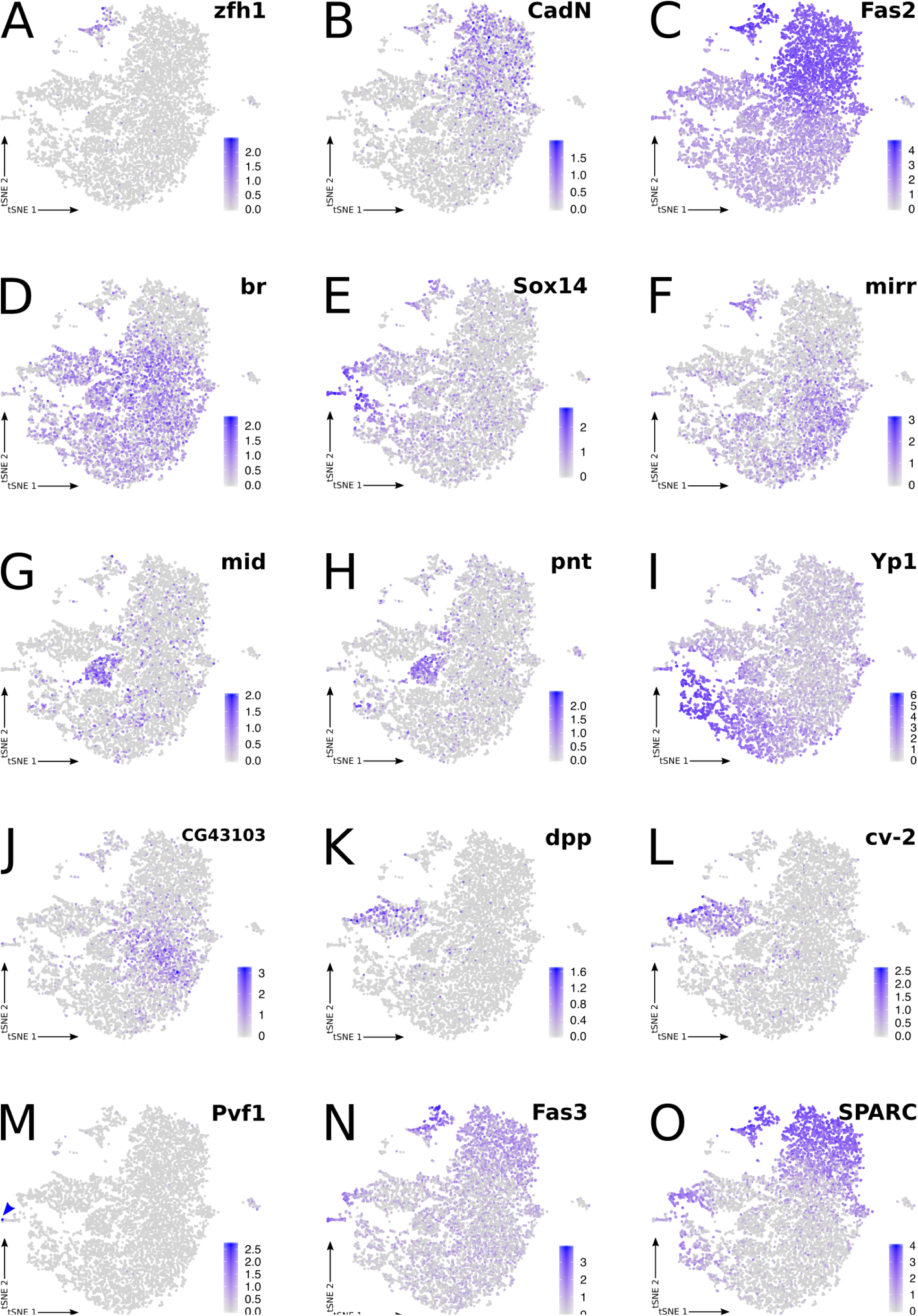
tSNE plots of selected genes. A-O) tSNE plots showing the expression pattern of the indicated genes.

**Figure S7:**
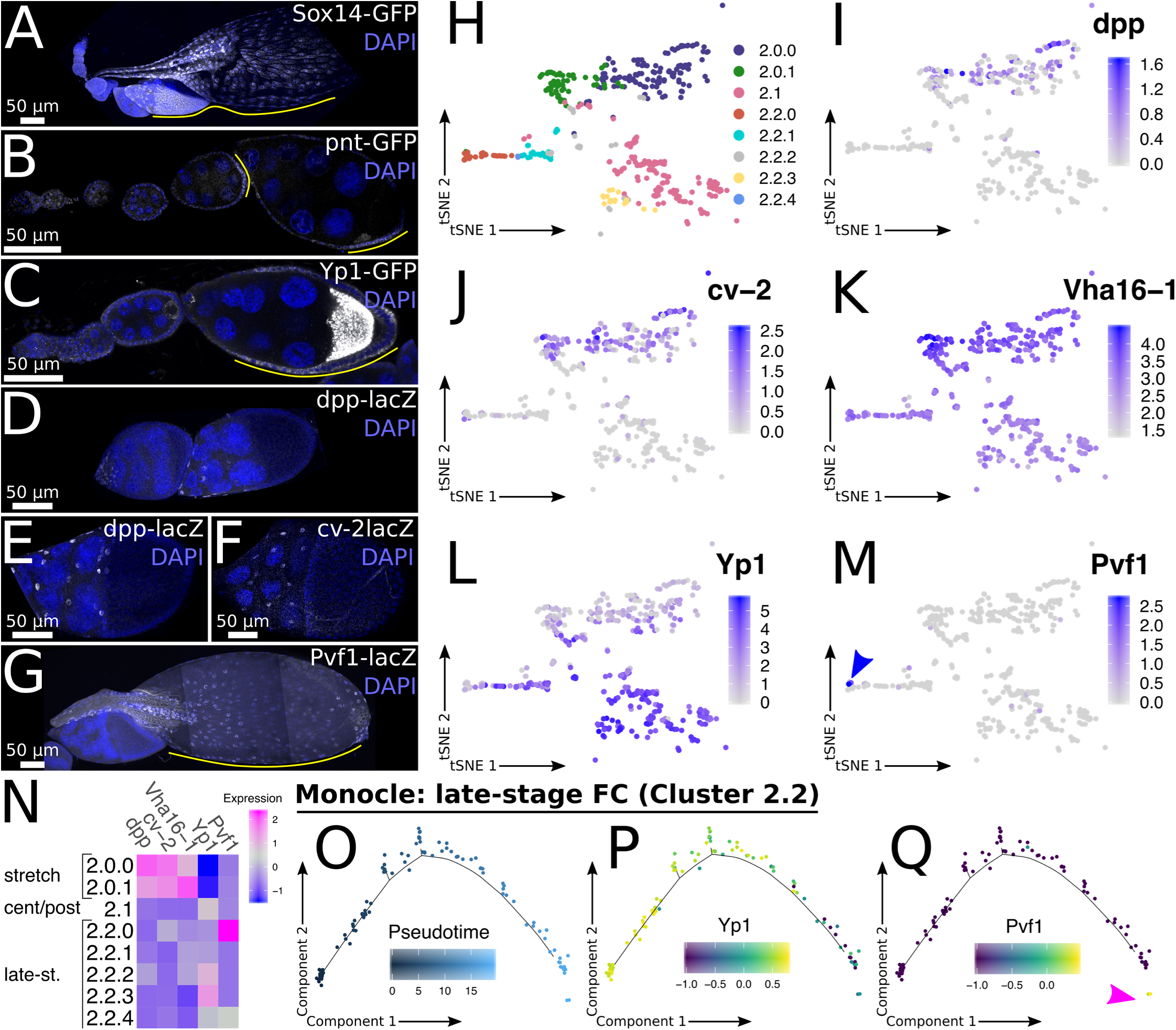
Markers of somatic cell populations in late stage follicles. A-G) Immunofluorescence staining of ovarioles. DAPI (blue) marks nuclei. Gene of interest in shown in white. Yellow line marks positive stages or cells. H) tSNE plot of Cluster 2. I-M) Expression of indicated gene on tSNE plot of Cluster 2. N) Heatmap of selected markers in Cluster 2. O-Q) Monocle analysis of Tier 2 Cluster 2.2 orders cells in pseudotime (O). Younger cells in 2.2 pseudotime express *Yp1* (P). The oldest cells in pseudotime express *Pvf1* (Q). Arrowheads point out positive cells for *Pvf1*.

**Figure S8:**
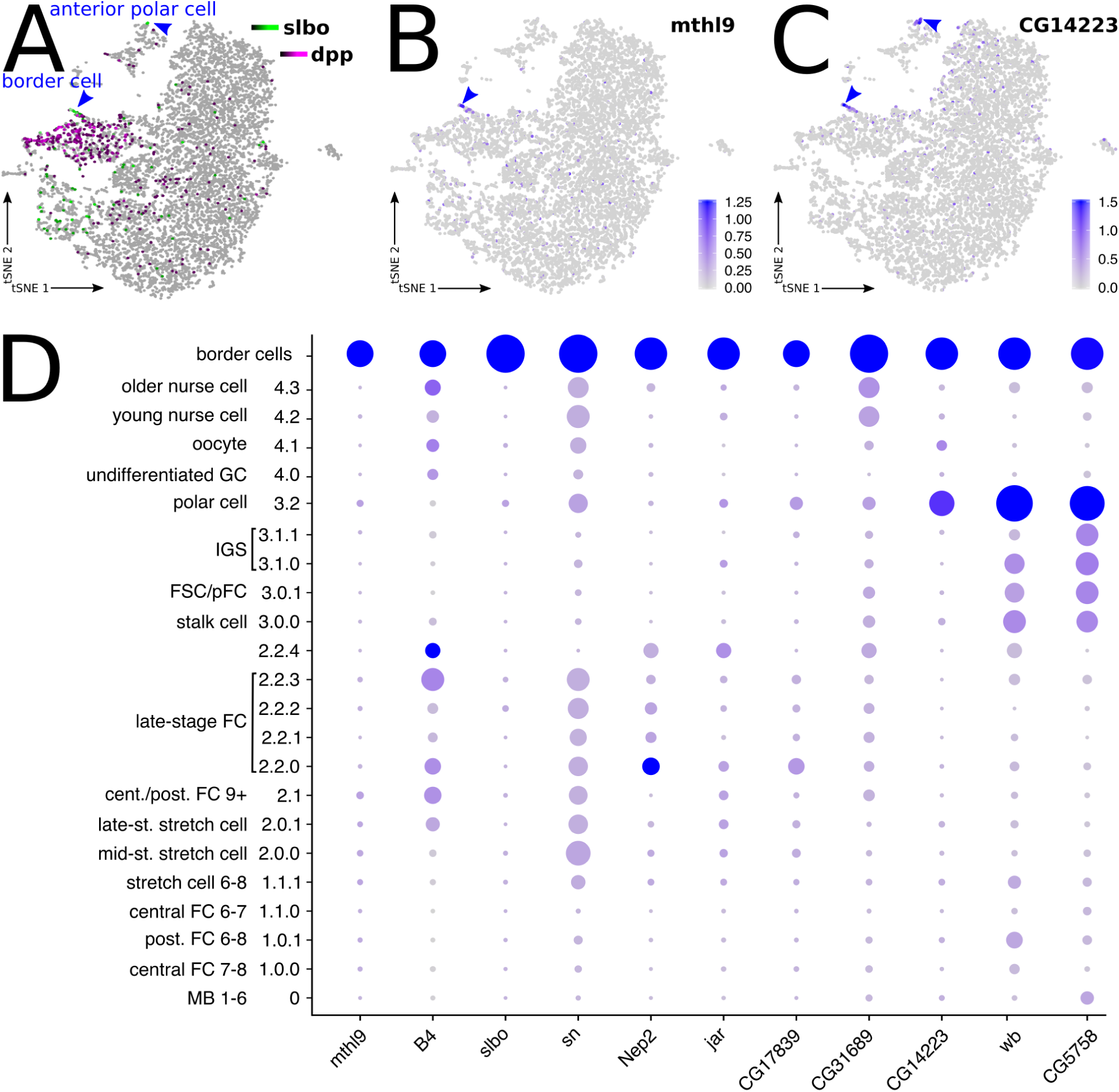
Transcriptional profile of border cells. A) SCope plot showing the expression of *slbo* and *dpp*. Border cells are identified as a subset of *dpp* positive cells in Cluster 1.1.1 which co-express *slbo*. *slbo* is also detectable in a subset of polar cells in Cluster 3.2, corresponding to anterior polar cells in stages with border cells. B-C) tSNE plots showing the expression pattern of the predicted border cell markers, *mthl9* (B) and *CG14223* (C). D) Dotplot showing the expression of border cell markers.

**Figure S9:**
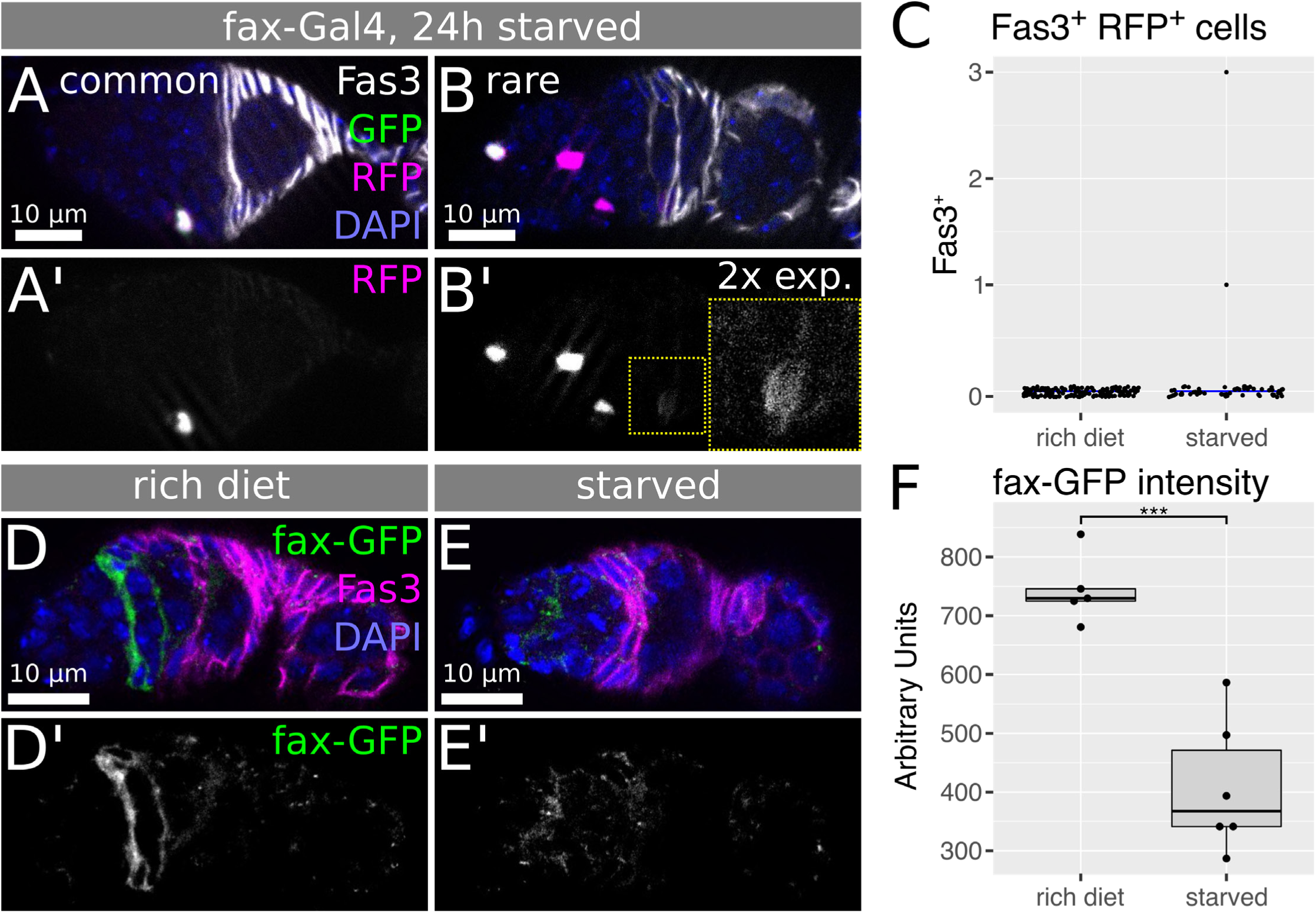
fax expression decreases in response to nutrient deprivation. A-B) Germaria with *fax-Gal4* driving G-TRACE^ts^ in starved flies immediately following a 24h exposure to the indicated diet condition stained for DAPI (blue), GFP (green), RFP (magenta), and Fas3 (white). A) Shows the commonly expression pattern, while B) shows very rarely observable RFP^+^ follicle cells with low signal. Inset shows region of interest with twice the pixel intensity setting that was used for the main image. C) Quantification of the frequency of Fas3*^+^* RFP^+^ cells. In flies fed a rich diet, RFP^+^ cells in the germarium are never Fas3*^+^*. In starved flies, RFP^+^ cells in the germarium are almost never Fas3*^+^* and, in the rare Fas3*^+^* RFP^+^ cell, shown in panel B, the RFP is very dim. n=165 for rich diet, n=126 for starvation. D-E) Immunofluorescence staining of fax-GFP germaria under rich diet (D) and upon starvation (E) stained and imaged with same settings. DAPI (blue) marks all nuclei. Fas3 (magenta) marks all follicle cells. Note that fax-GFP (green) expression is stronger when flies are kept on rich diet. F) Quantification of fax-GFP intensity in germaria cultured on rich diet or subjected to starvation. Intensity is significantly reduced upon starvation conditions. n=5 for rich diet, n=6 for starvation. Boxes in box plots show the median and interquartile range; lines show the range of values within 1.5x of the interquartile range. ***p = 2 x 10^-4^, using a two-sided Student’s T-test.

**Figure S10:**
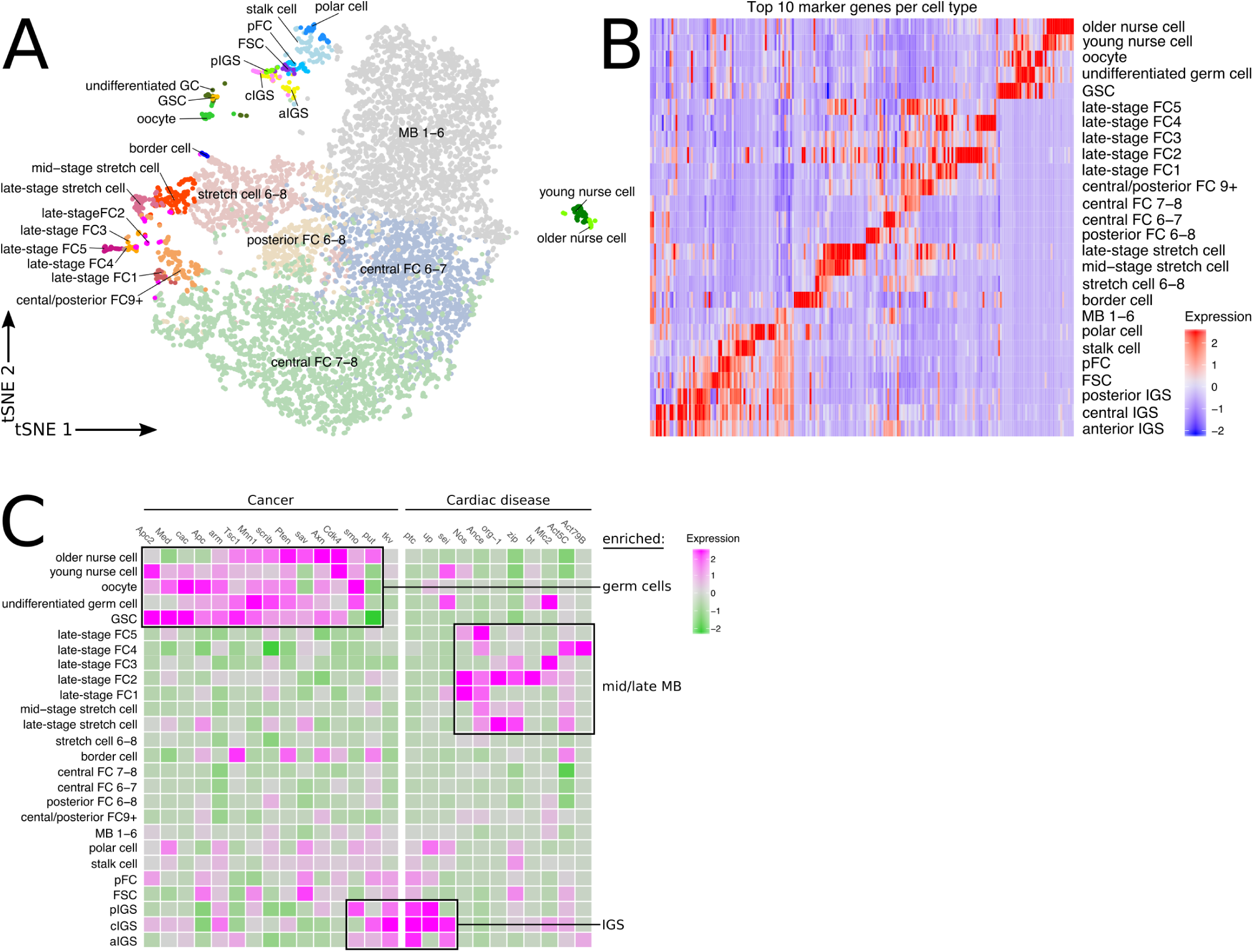
Drosophila ovarian cell types with human disease gene expression profiles. A) tSNE plot with all cell types identified in this publication. B) Heatmap showing expression of 10 markers per cluster across all identified cell types. C) Heatmap showing enrichment for cells expressing cancer-associated genes in germ cell clusters, cancer and cardiac disease-associated genes in IGS cell clusters, and cardiac disease-associated genes in main body follicle cell clusters.

